# Mapping cellular subpopulations within triple negative breast cancer tumors provides a tool for cancer sensitization to radiotherapy

**DOI:** 10.1101/2021.01.07.425553

**Authors:** Heba Alkhatib, Ariel M. Rubinstein, Swetha Vasudevan, Efrat Flashner-Abramson, Shira Stefansky, Solomon Oguche, Tamar Peretz-Yablonsky, Avital Granit, Zvika Granot, Ittai Ben-Porath, Kim Sheva, Amichay Meirovitz, Nataly Kravchenko-Balasha

## Abstract

Triple negative breast cancer (TNBC) is an aggressive type of cancer that is known to be resistant to radiotherapy (RT). Evidence is accumulating that is indicative of the plasticity of TNBC, where one cancer subtype switches to another in response to various treatments, including RT. In this study we aim to overcome tumor resistance by designing TNBC-sensitizing targeted therapies that exploit the plasticity occurring due to radiation exposure. Using single cell analysis of molecular changes occurring in irradiated TNBC tumors, we identified two initially undetected distinct subpopulations, represented by overexpressed Her2 and cMet, expanding post-RT and persisting in surviving tumors. Using murine cancer models and patient-derived TNBC tumors, we showed that only simultaneous targeting of Her2 and cMet was successful in sensitizing TNBC to RT and preventing its regrowth. The strategy presented herein holds the potential to be broadly applicable in clinical use.

**Highlights:**

- Sensitization of TNBC to radiotherapy (RT) is a clinically unmet need
- Single cell strategy creates a precise map of subpopulations expanding post-RT
- Evolution of intra-tumor heterogeneity is turned into a therapeutic advantage
- Simultaneous targeting of expanding subpopulations sensitizes TNBC to radiotherapy

## Introduction

Radiotherapy emerged more than a century ago^1^, and continues to be a key modality in the treatment and management of various types of cancer.

Recent studies have shown that radiation, while effectively kills cancer cells, also promotes anti-apoptotic and pro-proliferative responses that often result in tumor regrowth^1,2^. This notion gave rise to numerous studies attempting to characterize tumor molecular phenotypes occurring in response to radiation, in order to develop new strategies to enhance the response of cancer to radiotherapy (see for example^3–8^).

Triple negative breast cancer (TNBC) is a clinically unique, aggressive and highly heterogeneous subtype of breast cancer that does not express estrogen receptors, progesterone receptors or human epidermal growth factor receptor-2 (Her2), and for which not a single targeted therapy has been approved. Chemotherapy (CT) and radiation therapy (RT) have therefore remained the standard treatments over the past 20 years^9,10^. Although TNBCs can be sensitive to RT during early treatment stages, they often develop resistance at later stages^10^. One of the main reasons for this is significant variability between tumor cells within the tumor, which poses a major obstacle in the treatment of TNBC tumors^11,12^. Moreover, accumulating evidence for the plasticity of breast cancer cells, such as a switch from Her2^-^ to Her2^+^ phenotypes in response to RT^13^, may complicate breast tumor classification and thus the design of appropriate therapy.

Finding a strategy with the ability to transform the potential evolution of certain subpopulations within irradiated TNBC tumors into a therapeutic advantage, is an unmet need in cancer research and clinical practice.

We propose a novel concept according to which TNBC sensitization can be rationally-designed based on the resolution of patient-specific intra-tumor subpopulations and examination of the dynamics thereof in response to RT treatment.

Considering that even very small subpopulations within the tumor may eventually give rise to tumor regrowth (e.g. tumor stem cells), it is vital that the identification of tumor subpopulations will be of high resolution.

Herein we employ an information theoretic technique, surprisal analysis (SA), to resolve TNBC cellular subpopulations on the single cell level, evaluate their response to RT, and design a therapeutic method to sensitize TNBC cells to RT.

We consider tumors to be homeostatically disturbed entities, which have deviated from their balanced state due to various constraints (e.g. mutational stress, application of drugs, etc.) ^14^. Each constraint creates a deviation in the expression levels of a subset of proteins in the tumor. Thus, a constraint creates an <u>unbalanced process</u> in the tumor, consisting of the group of proteins that were altered by the constraint. SA examines protein-protein correlations and, based on information theoretic and thermodynamic-like considerations, identifies the constraints that operate in the studied system as well as the proteins affected by each constraint.

We have previously demonstrated that the accurate identification of the signaling network structure emerging in MCF10a human mammary cells upon stimulation with EGF, allowed us to anticipate the effect of the addition of protein inhibitors on the protein network structure^15^. In other studies we have shown that SA of cell-cell signaling in brain tumors provides a predicition about cellular spatial distributions and the direction of cell-cell movement^16^. We have also implemented SA in large-scale proteomic datasets obtained from multiple cancer types and demonstrated how this analysis successfully predicts efficient patient-specific targeted combination therapies^17^.

In this research, we utilize SA to study single cells. For each cell we identify a cell-specific signaling signature – **CSSS**, consisting of a set of unbalanced processes that have emerged within the individual cell. We then define an intra-tumor subpopulation to be a group of cells harboring the exact same CSSS. These cells are expected to respond similarly to treatment. The final result of the analysis is a high-resolution intra-tumoral map of the different subpopulations within the tumor, and the CSSS that operates in every subpopulation. Importantly, even very small subpopulations comprising less than 1% of the total detected population can be captured using our strategy. Such a robust and comprehensive map has the ability to provide guidance on the accurate determination of drug combinations to effectively target dominant subpopulations, as well as small and persistent subpopulations within the tumor, and bring about a potent therapeutic effect.

A 4T1 murine model for stage IV TNBC^18^ is utilized in this study, as well as human TNBC and patient-derived xenograft models. We show that upon RT treatment in-vitro and in-vivo, all models demonstrate a significant expansion of two distinct cellular subpopulations: one with upregulated EGFR/Her2 and another with upregulated cMet/MUC1. These subpopulations are barely detectable in untreated tumors. We believe that the poor response of TNBC to RT can be overcome by inhibiting the growth of these subpopulations. We validate our hypothesis by showing that RT-treated TNBC tumors that are simultaneously pretreated with both anti-Her2 (Trastuzumab) and anti-cMet (Crizotinib) inhibitors do not relapse in-vitro or in-vivo. Assessment of each targeted drug alone demonstrates a significantly smaller effect.

In summary, this study provides a novel framework for the resolution of tumor-specific cellular heterogeneity at the single cell level. We show that accurate mapping of tumor cellular subpopulations within a TNBC mass can provide guidance on how to incorporate targeted therapy with RT in order to overcome resistance.

The proposed approach is expected to augment the success of radiotherapy in clinical oncology and significantly improve the outcome of TNBC and potentially other cancer types.

## Results

### Overview of the integrated experimental-computational approach

We based our study on the notion that TNBC tumors that undergo irradiation treatment, while initially responding to the treatment, eventually relapse and regrow (**Fig. 1**). We hypothesized that the ability of the tumors to relapse stems from the existence of intra-tumoral subpopulations that do not respond well to the irradiation treatment and drive the regrowth of the tumor post-radiation (**Fig. 1**, top).

**Figure 1.**
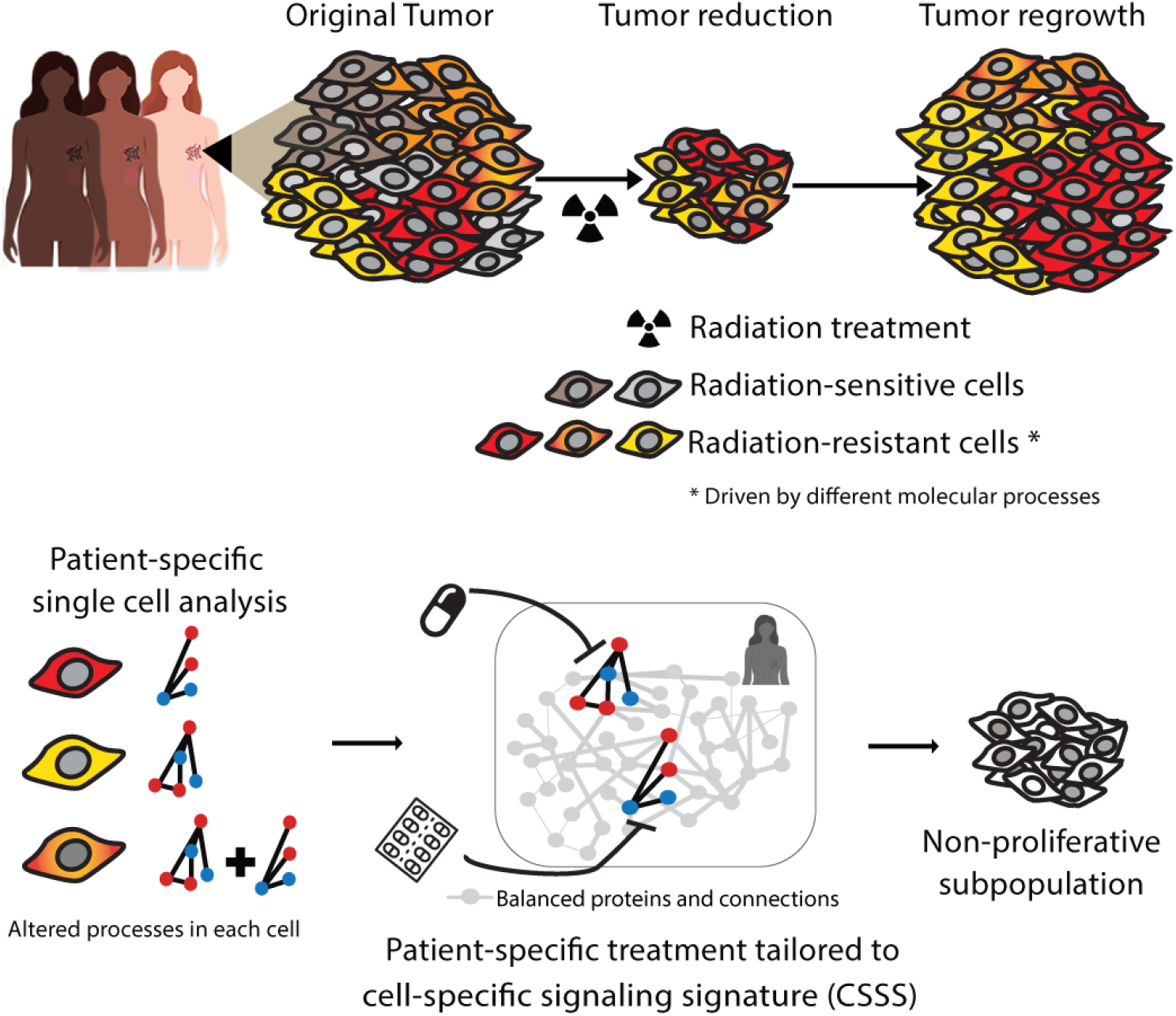
Decoding intra-tumor heterogeneity into distinct subpopulations after radiation treatment offers potential new targets for tumor-specific therapy. Top: Phenotypic variations due to intra-tumor heterogeneity pose a significant challenge in attaining optimal patient-specific therapy regimens. Bottom: Utilizing high throughput flow cytometry and single cell surprisal analysis, patient-specific tumor network structures are identified, with the elucidation of cellular subpopulations and altered processes in each subpopulation, pre- and post-radiotherapy. Accurate targeting of resistant subpopulations aims to prevent cellular expansion by sensitizing the tumor to RT.

We set out to study TNBC tumor composition on the single cell level, aiming to identify a set of intra-tumoral subpopulations, including relatively small subpopulations, that exhibit a diminished response to RT treatment. By elucidating the altered molecular processes that each subpopulation harbors, we devised a therapeutic strategy believed to intensify the response of the tumor to RT (**Fig. 1**, bottom).

Several dimensionality reduction algorithms have been developed to interpret single cell variations (e.g. variations in protein or gene expression levels), such as clustering-based t-SNE analysis^19^ or principle component analysis (PCA)^20–22^. These methods are very useful in statistical determination of dominant expression patterns but are limited when a more deterministic partitioning of the tumor mass into cellular subpopulations, based on cell-specific sets of altered molecular processes, is required. For example, t-SNE is a non-deterministic method (e.g. different runs with the same hyper parameters may produce different results) and is unable to assign a certain protein to several processes, or to determine which processes are active in every cell. Therefore t-SNE will be less efficient when the determination of robust cell-specific signaling signatures is required (e.g. for drug combination design). Similarly, PCA focuses mainly on the most dominant patterns obtained from proteins with the highest variability in the population, rather than on cell-specific sets of altered processes (for more details see references ^23,24^).

We sought a deterministic approach, in which every single cell can be plotted according to its molecular aberrations and network reorganization. To this end, we employed surprisal analysis (SA), an information theoretic method^14,25^ originally applied to characterize the dynamics of non-equilibrium systems in chemistry and physics^26,27^. This analysis has recently been utilized to quantify bulk proteomic changes in large datasets, including multiple patient tissues and cancer cell lines, in order to predict a change in the behavior of the systems^15,28^ or design individualized drug therapies^17^.

Herein, we extend the approach to quantify expression changes in single cells in order to accurately characterize the changes occurring in tumor cellular populations in response to RT. Our analysis is based on the premise that the application of radiotherapy to TNBC cells induces certain constraints within the tumor mass. These constraints result in altered expression levels of certain proteins relative to their balanced levels. This in turn reflects the plasticity of the tumor in response to RT. SA recognizes the constraints operating in the system by identifying groups of proteins that exhibit similar deviations from their balanced state (**Fig. 1**). A group of proteins demonstrating similar alterations in expression patterns is defined as an *unbalanced process*. Hence, every constraint that operates on the system gives rise to an unbalanced process. SA identifies the unbalanced processes that operate in the system under study, including the group of proteins affected by each process. Each protein may participate in several processes ^17^. It is important to note that not all processes are active in all cells, i.e. a certain process can have negligible amplitude in some cells and significant amplitude in others. A number of different unbalanced processes may operate simultaneously in every cell (**Fig. 1**). The cell-specific signaling signature (CSSS) is defined for each cell, according to the set of active unbalanced processes in that specific cell (See Methods for additional details).

To collect high resolution data regarding the intra-tumoral composition of TNBC tumors, we employed the following experimental technique: Samples obtained from multiple sources (e.g. cell lines, mouse models and patient-derived tumor cells) were processed into single cell suspensions. The cell suspensions were then labeled with fluorescently-labeled antibodies targeting selected cell-surface oncoproteins and assayed by multicolor FACS to reveal accurate protein expression levels in each single cell.

In each experimental condition approximately 30,000-50,000 single cells were profiled allowing for the identification of different subpopulations, including very small subpopulations (comprising less than 1% of the total population) that have significantly limited detection rates when using standard pathological tests.

#### Selection of oncomarkers for single cell analysis

The selection of the protein panel for FACS analysis was based on unbiased gene expression data analysis of TNBC tumors responding to cytotoxic stress (irradiation and chemotherapy) and an extensive literature search to filter out oncoproteins that best represent possible expression patterns in TNBC cells^29–35^.

Gene expression data was obtained from 14 patient-derived TNBC tissues that were irradiated and treated with Taxol^36^. Four of the patients achieved pathologic complete response (pCR), while 10 remained with residual disease and were classified as non-pCR (NR, **Fig**.**2a**). First, using unbiased *bulk* surprisal analysis, we identified co-expressed altered gene expression patterns, *unbalanced processes*, characterizing the variability of the dataset ^14,24^ (**Fig**.**2a**). The dataset (28 samples-representing 14 tumors measured in duplicate) was characterized by 14 unbalanced processes (**Fig**.**2b**, Methods). To provide biological interpretation of these processes, transcipts with significant weights (**Fig**.**2a**, Methods and Table S1) were classified into biological categories based on Gene Ontology (Methods, Table S1). For example, process 1+ included upregulated transcripts involved in multiple categories belonging to the cell cycle and signal transduction, while process 2+ to the cell cycle and DNA damage (Table S1). Biological characterization of all processes is provided in Table S1. Central transcripts that could serve as representatives of these 14 processes were seeked next.

Eleven cell-surface oncoproteins capturing the gene expression variability in TNBC patients were selected, and appeared both in pCR and non-pCR patients: Her2, EGFR, EpCAM, CD44, CD24, PD-L1, KIT, CD133, E-Cadherin, cMet and MUC1 (**Fig. 2c**). These oncomarkers are also known to be involved in breast cancer/cancer stem cell proliferation and represent potential drug targets for therapy or biomarkers for diagnostics^29–35,38^. To examine the response of TNBC to irradiation, single cell measurements were performed in which the selected 11 biomarkers were quantified in each cell.

**Figure 2.**
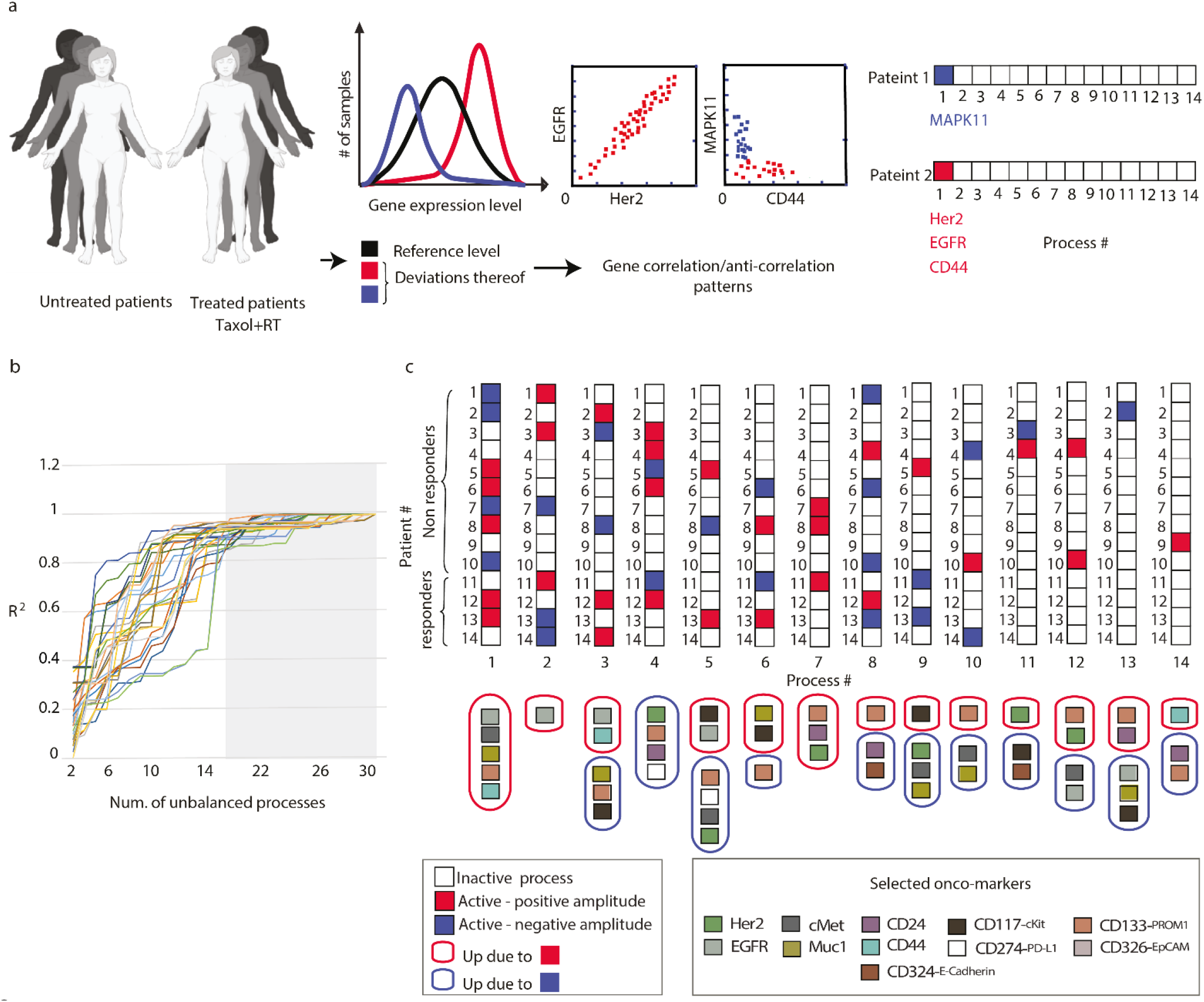
Utilizing gene expression profiling to select oncomarkers for single cell analysis. (**a**) Quantitative gene expression data obtained from a cohort of TNBC tumors irradiated and treated with Taxol, is used for surprisal analysis. Extent of variability in gene expression levels is quantified for each transcript, and further visualized and quantified using gene expression distribution histograms. Transcripts whose expression levels deviate from the reference state in the same or opposite directions, i.e. co-varying transcripts, are grouped further into correlation networks (exemplified here by EGFR, HER2, MAPK11 and CD44). In this example EGFR and HER2 are correlated, whereas the expression levels of MAPK11 and CD44 are anticorrelated and deviate from the steady state in an opposite manner. (**b**) 14 significant unbalanced processes (Table S2) were found in the dataset based on error calculation that characterize gene expression variability in the dataset^24,37^. (**c**) A patient-specific combination of unbalanced processes was calculated for every patient (see also Table S2). Combinations were generated using amplitude values that exceeded threshold limits calculated as explained previously^24^. For example patient 1 has three active processes: 1, 2 and 8. Negative/positive amplitude denote how the patients are correlated with respect to a particular process. 11 differentially expressed transcripts (Her2, EGFR etc.,lower panel), belonging to various biological categories, such as cell proliferation, motility, EMT and cancer stem cells, were found to participate in different 14 processes and were selected further as representatives of this variability.

#### Single cell SA

FACS measurements (**Fig. 3a**) were analyzed by single cell SA to reveal proteins that demonstrate deviations in expression levels relative to their reference state levels (**Fig. 3b**). Cell-specific protein-protein correlation expression patterns were then examined (**Fig. 3c,d**) to identify newly emerged unbalanced processes as well as the sets of unbalanced processes that operate in specific cells, namely the **CSSS** (**Fig. 3d**). Each CSSS is graphically represented by a cell-specific barcode where white squares indicate inactive and black/gray indicate active processes in a cell (**Fig. 3d**, right panel). We then define cellular subpopulations as groups of cells harboring the same CSSS (**Fig. 3e**).

**Figure 3.**
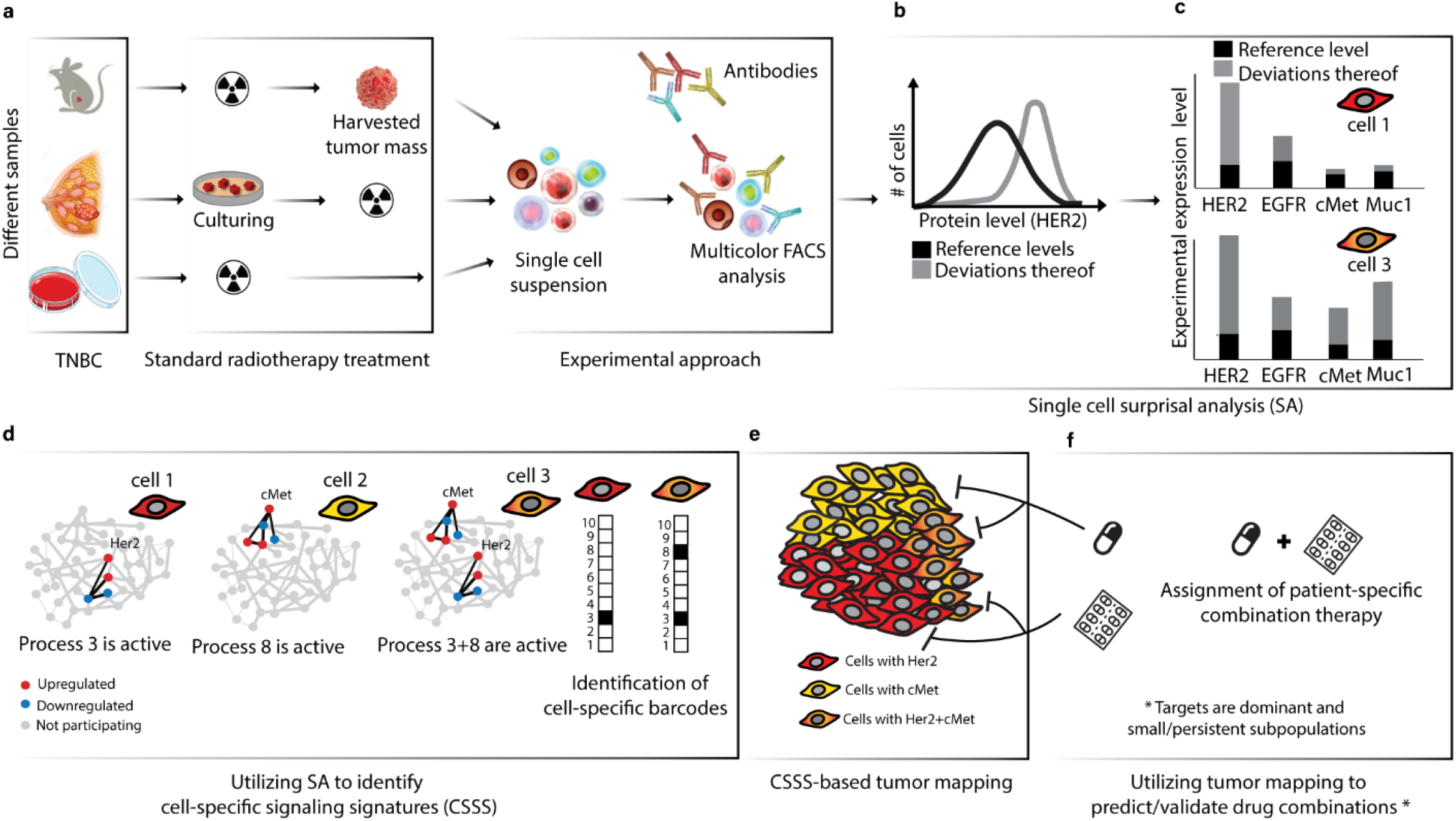
Schematic of the application of the surprisal analysis algorithm. (**a**) Preparation of fluorescently-tagged single cell suspensions from different sample sources post-irradiation for cytometry analysis (right panel). **(b)** Using 30,000-50,000 cells/sample, surprisal analysis reveals protein expression level distributions at the reference (balanced) state and deviations thereof. (**c**) Proteins that deviate in a similar manner from the references (e.g. both induced in a certain group of cells) are grouped into altered subnetworks, referred to as “unabalanced processes”. (**d**) Unbalanced processes with significant amplitudes are assigned to each cell providing a cell-specific signaling signature (CSSS). Each CSSS is transformed into a cell specific barcode (right panel). (**e**) Cells sharing the same barcode are organized into distinct subpopulations. (**f**) Tumor-specific targeted therapy combinations are tailored to the CSSS.

Note that different subpopulations may share similar processes, e.g. the red and orange subpopulations in Figure 3 both harbor unbalanced process 3 (**Fig. 3d**). However, the <u>complete set</u> of unbalanced processes in each subpopualtion, namely the CSSS, is what governs the therapeutic strategy that should be taken.

The in-depth information collected in the previous steps is utilized to devise a therapeutic strategy that incorporates targeted therapies to aid RT. This is achieved by targeting the dominant and RT-resistant subpopulations, to potentially achieve long term tumor remission (**Fig. 3f**).

### 10 unbalanced processes give rise to the expression variations of 11 cell-surface proteins in 4T1 mouse TNBC cells

4T1 cells, obtained from a spontaneously developed tumor in an immunocompetent murine model for stage IV TNBC^18^, were irradiated using two doses (5 Gy or 15 Gy), and then grown under standard conditions for 24h, 48h and 6 days. The cells were then suspended and the expression levels of the selected panel of 11 cell-surface oncoproteins in single cells were measured using FACS. Figures **4a** and **S1** show the overall distributions of expression levels of the different proteins in the cells measured.

Two-dimensional correlation plots were created to gain insight into the behavior of these proteins in single cells. For example, although cMet and Her2 were both upregulated in response to RT, their expression levels showed poor correlation (**Fig. 4b**), where on the contrary, EGFR and Her2 levels demonstrated a strong correlation (**Fig. 4c**), as did MUC1 and cMet expression levels (**Fig. 4d**).

**Figure 4.**
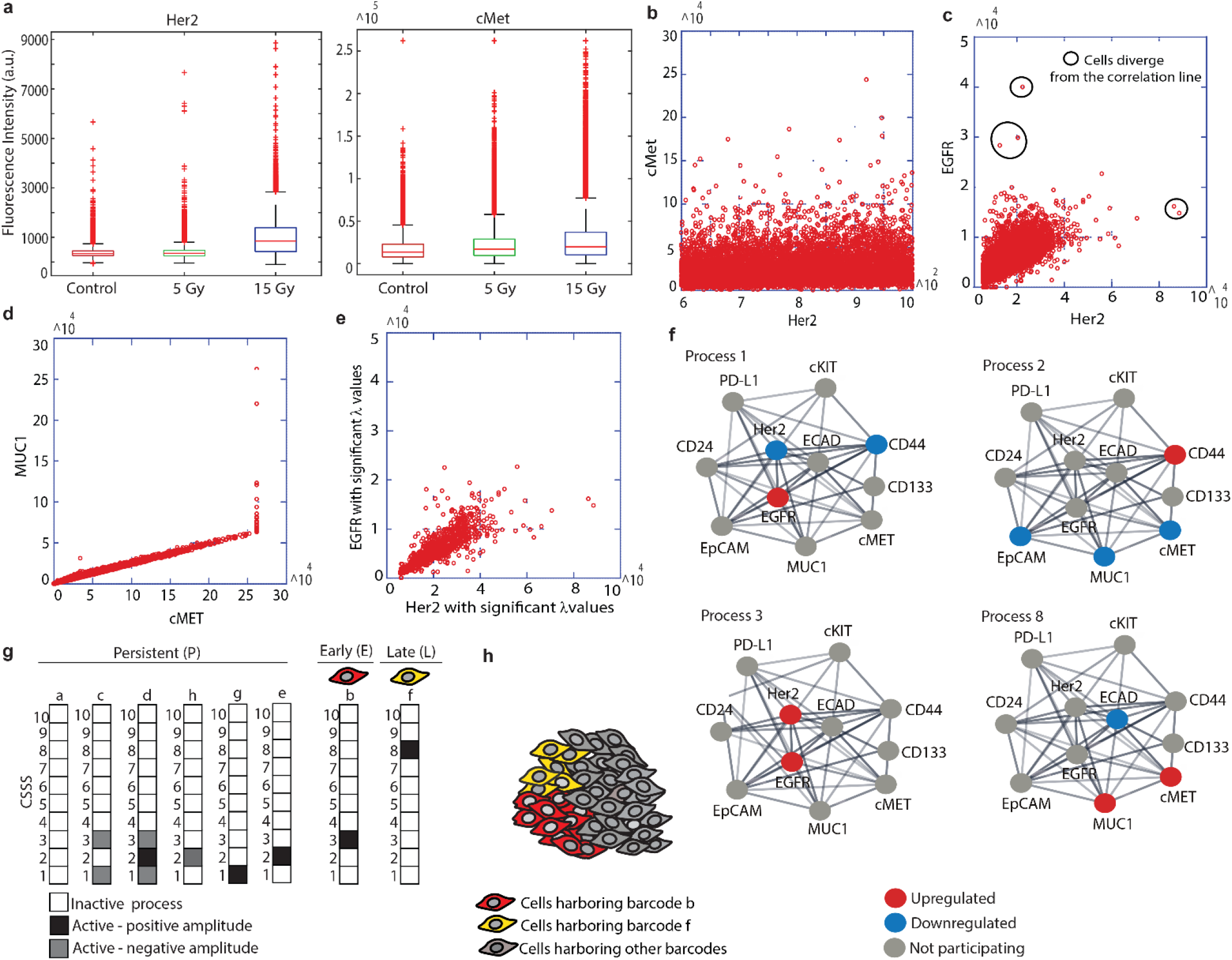
Resolution of expanded subpopulations in 4T1 cellular population post irradiation. (**a**) FACS expression levels of Her2 and cMet following RT. (**b-d**) FACS experimental data plotted as correlation plots between Her2 and cMet (b), Her2 and EGFR (c), and MUC1and cMet (**d**). (**e**) Correlation plot between Her2 and EGFR expression levels including the cells with significant *λ*_3_ (cell) values (Methods). (**f**) Four examples of 10 unbalanced subnetworks (processes) resolved in 4T1 are shown. Protein-protein interactions were determined using the String database. (**g**) Each cell was assigned a barcode representing CSSS. Most abundant (>1%) subpopulations are presented. Based on these CSSSs the tumor was divided into distinct subpopulations (**h**).

Note, however, that a strong correlation between EGFR and Her2 does not necessarily mean that these proteins participate in the same unbalanced processes in all tested cells. Small subpopulations of cells unaffected by the same processes, and possibly displaying a poorer correlation between these proteins, may be overlooked when studying variations in all cells simultaneously (**Fig. 4c**, black circles). Similarly, small subpopulations of cells that demonstrate a <u>strong</u> correlation between cMet and Her2 may exist, but nevertheless be masked by the representation shown in Figure 3b. Moreover, the expression level of a certain protein can be influenced by several processes due to non-linearity of biochemical processes: a certain pair of proteins can be correlated or non correlated in the different unbalanced processes operating in the same cell, further complicating the interpretation of these 2D correlation plots. We therefore performed single cell SA to map the unbalanced processes operating in the entire cellular population as well as in each single cell (see Methods and Poovathingal et al.^25^ for details). The analysis revealed 10 unbalanced processes (i.e. altered protein-protein correlation patterns resulting from 10 constraints) occurring in the untreated/treated cells (**Fig. 4f, Fig. S2**). Four of the processes, all appearing in at least 1% of the treated and/or untreated cells are shown (**Fig. 4f**; the remaining unbalanced processes are presented in **Fig. S2**). The most abundant processes, indexed 1 and 2, appeared in 25% and 18% of the untreated cells, respectively. Processes 3 and 8, which included correlated Her2/EGFR and cMet/Muc1, respectively, initially demonstrated low abundancy, and appeared in 0.3% and 0.5% of the untreated cells, respectively (**Fig. 4f, Table S3**). Processes 3 and 8 became more dominant 6 days post-RT (**Table S3**; more details in the next sections).

### 8 sets of unbalanced processes, or 8 distinct CSSS’s, were resolved suggesting that the cells form 8 distinct subpopulations

As mentioned above, more than one unbalanced process can operate in every cell. Therefore, to gain in-depth information regarding the complete altered signaling signature in each cell, we examined the <u>sets</u> of unbalanced processes in the cells studied, namely the CSSS.

We found that 8 different sets of unbalanced processes, representing 8 distinct signaling signatures (CSSS), repeated themselves in the population of cells before and/or after RT (**Fig. 4g**). For the simplicity of representation, each CSSS was translated into a cell-specific barcode in which active/inactive processes were color-labeled (**Fig. 4g**).

Only unbalanced processes with significant amplitudes were included in the CSSS of each individual cell (**Fig. S3, S4** and Methods). Figure **3e** shows how selecting only cells with high amplitudes improves the correlation between relevant proteins within the processes and thus the accuracy of the unbalanced processes in the analysis.

Additionally, only CSSS’s that appeared in at least 1% of the cells were taken into account. The barcodes of these abundant subpopulations consisted of processes 1, 2, 3 and 8 (**Fig. 4f, g**).

### The 8 abundant cellular subpopulations demonstrate different te5mporal behaviors and different variations in abundance

When we examined the temporal behavior of the abundant subpopulations, we found that they could be divided into 3 groups: (1) <u>persistent</u> subpopulations, which decreased 48h post-RT and then returned to their previous size 6d post-RT; (2) <u>early</u> subpopualtion, which was minor initially (<1%), and then expanded 48h and 6d post-RT (>1%); (3) <u>late</u> subpopulations, which was small prior to RT and expanded 6d post-RT (a schematic representation of the different temporal behaviors is shown in **Fig. 5a**).

**Figure 5.**
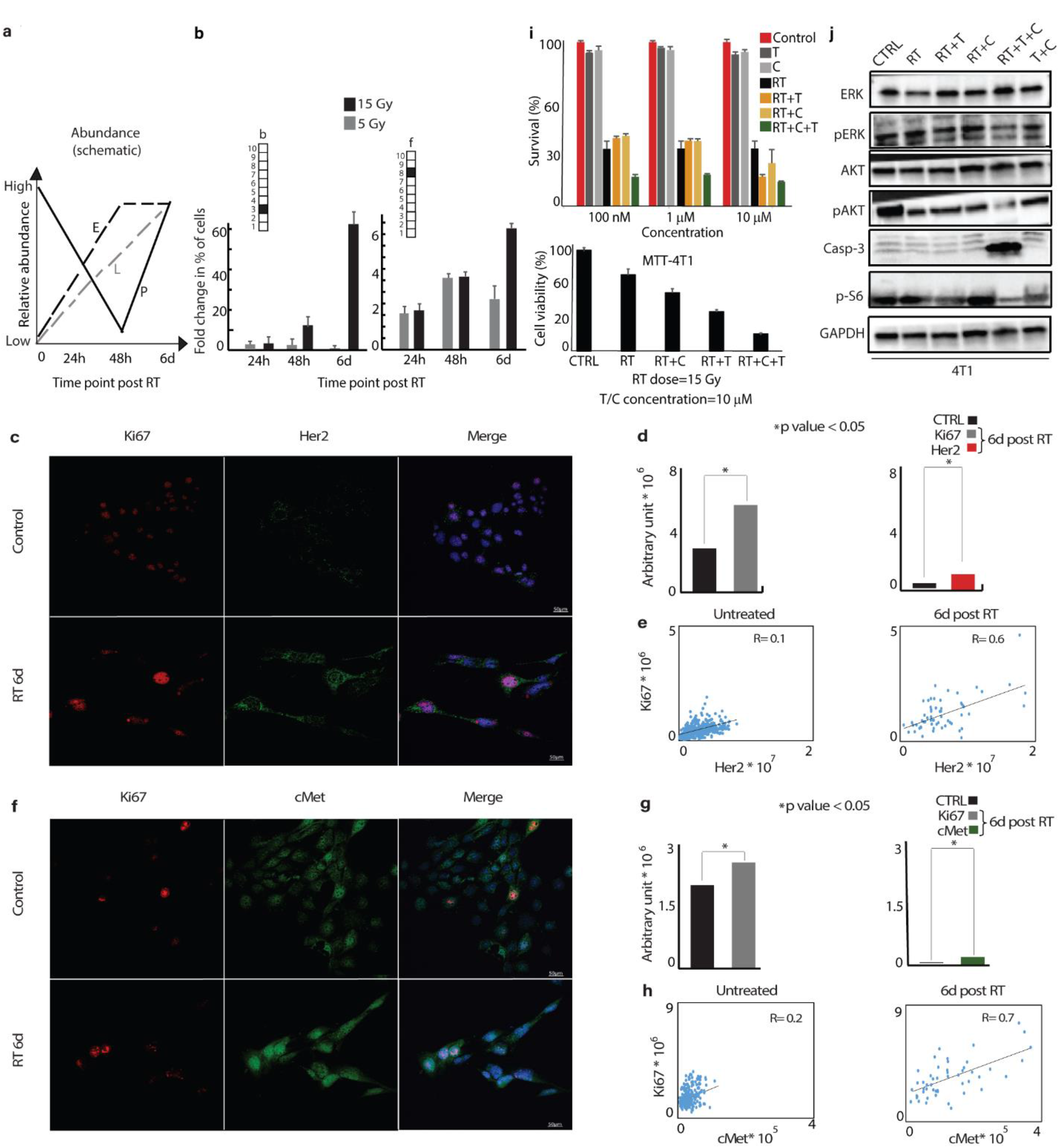
Two distinct subpopulations expand and show proliferative properties in response to RT. **(a)** Schematic representation of the temporal behavior of abundant subpopulations is demonstrated. **(b)** Very small subpopulations (<<1%), represented by barcodes ***b*** and ***f***, expanded significantly following RT. **(c-h)** 4T1 cells were irradiated with 15Gy. 6 days post-RT, cells were incubated with antibodies against Ki67, cMet and Her2 and nuclei were stained with DAPI (fluorogel II)(**e**,**h**) 40x lens; scale bar represents 50µm. (**f**,**i**) Sum intensities of Ki67 (f, left panel); Her2 (f, right panel); Ki67 (i, left panel); cMet (i, right panel) were calculated from 8-10 fields using the NIS-Elements software (Nikon); *P-value < 0.05. (**g**,**j**) Correlation plots between Ki67 and Her2 (**g**) and Ki67 and cMet (**j**) were generated for each indicated condition to test co-activation represented in **(e**,**h)**. R values indicating the extent of correlation between Ki67 and Her2 (g) Ki67 and cMet (j) were calculated before and after RT. (i) Survival rates of 4T1 cells in response to Trastuzumab (T), Crizotinib (C), RT, RT+C, RT+T and RT+T+C as detected by MB survival assays 6 days post RT (upper panel), and cell viability as measured by MTT assay (lower panel). Drugs were added from 3 days prior to RT until the end of the experiment. SD are shown. **(j)** Downstream Her2 and cMet signaling, as represented by key downstream proteins, is shown following different treatments. Our predicted combination induced high levels of cleaved caspase-3 compared to radiation alone, irradiation+**T** and irradiation+**C**. Downregulation of pAKT, pERK and p-S6 was detected when **T**+**C** was applied prior to RT.

The abundance of persistent subpopulations did not change 6 days post-RT treatment. For example, subpopulation **c** comprised 14.5% of the cells before RT, and a similar percentage of the cells, 14.2%, was found to comprise this subpopulation 6 days post-RT. However, early and late subpopulations, **b** and **f**, respectively, expanded significantly 6 days post-RT (**Fig. 5b**). Subpopulation **b** harbored only process 3, in which Her2 and to a lesser extent EGFR (**Fig**.**4g**), were induced (**Fig. 5b, Table S3**). Strikingly, subpopulation **b** was induced 60-fold post-irradiation relative to the non-irradiated cells (expanded from low (<1%) levels in untreated cells to ~19-22% of the population, 6 days post-RT, **Fig. 5b**).

Subpopulation **f** harbored only process 8 (**Fig. 4g**), with induced cMet/MUC1 and reduced ECAD. Significant induction of subpopulation **f** was also observed, from undetectable levels to ~4% 6 days post-RT (**Fig. 5b)**.

These results demonstrate an important concept: although cMet and Her2 were both induced in response to RT (**Fig. 4a**), CSSS-based analysis revealed that those two proteins were expressed in *distinct* cellular subpopulations (processes 3 and 8 do not appear in the same cells; **Fig. 4g**). The development of such large, distinct and well-defined Her2+ and cMet+ subpopulations post-RT suggests that Her2 and cMet signaling may play a significant role in 4T1 cell survival and resistance in response to irradiation.

### Her2 and cMet positive subpopulations demonstrate proliferative properties

To characterize proliferative properties of the expanded Her2+ and cMet+ subpopulations in response to RT, we co-stained the 4T1 cell population with anti-Ki67 (proliferative biomarker) and anti-cMet and HER2 antibodies using immunofluorescent assays. Ki67, Her2 and cMet expression increased significantly in the cells surviving RT (**Fig. 5c,d,f and g**). Moreover, this result was supported by enhanced coordinated expression of Her2 and Ki67 (**Fig. 5e**) as well as cMet and Ki67 (**Fig. 5h**) proteins respectively as represented by an increased correlation between Her2 and Ki67; and cMet and Ki67 proteins, post-RT. This enhanced correlation in protein expression reveals the increased proliferative properties of Her2 or cMet expressing cells.

### Simultaneous inhibition of Her2 and cMet sensitized 4T1 cells to RT treatment

We hypothesized that simultaneous inhibition of both proteins, and thus targeting of both subpopulations, may sensitize 4T1 cells to RT. Her2 and cMet represent good candidates for such a strategy, as they are both druggable oncoproteins against which FDA-approved drugs exist.

To validate this hypothesis, we inhibited either each protein alone or in combination, beginning 2 days prior to RT and until 6 days post-RT, afterwhich cell survival was measured.

The Her2 inhibitor, Trastuzumab **(T)** and cMet inhibitor, Crizotinib **(C**), showed a synergistic effect in sensitizing the cells to RT (**Fig. 5i**). The combination of both drugs with RT increased cell death and also brought about depletion of signaling downstream to Her2 and cMet, as indicated by the low levels of downstream signaling proteins pERK1, pAkt and pS6K and the enhanced cleavage of the apoptotic marker Casp3 (**Fig. 5j)**.

### Her2+ and cMet+ cellular subpopulations expanded in response to RT in-vivo

To validate our hypothesis further, we implanted 4T1 cells into Balb/c mice, an immunocompetent murine model for TNBC. The cells were irradiated post-implantation using brachytherapy-focused irradiation technology adapted for mice^39^ by CT imaging and Monte-Carlo based dosimetry (**Fig. 6a**). 4T1 tumors were then isolated and single cell suspensions were analyzed.

**Figure 6.**
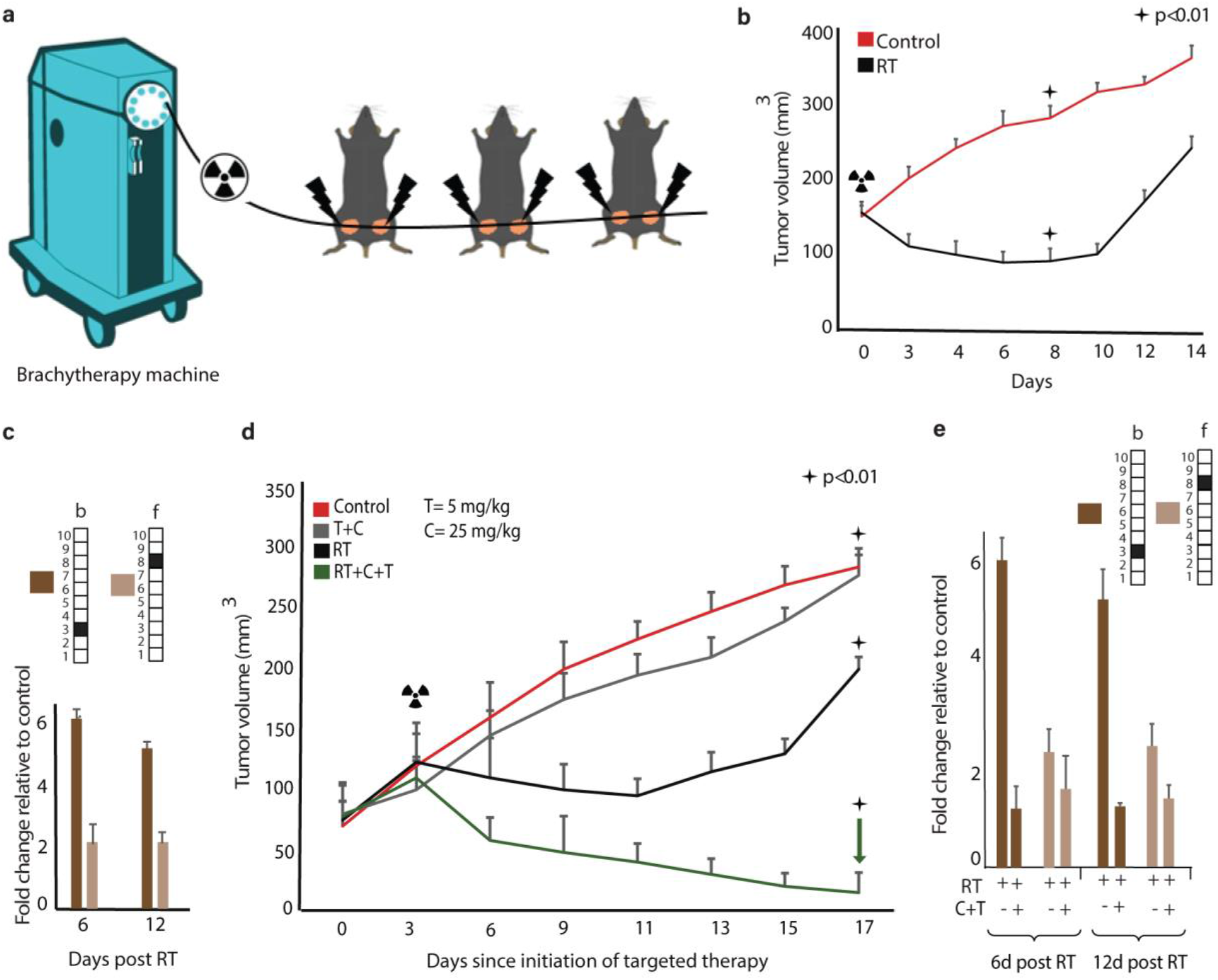
Inhibition of RT-induced subpopulations sensitized tumor response to RT. **(a)** 6-7 week-old Balb/C female mice were subcutaneously injected with 4T1 cells. When tumor volumes reached 80-100 mm^3^, mice were treated with brachytherapy RT on alternate days (12 Gy). **(b)** Tumor volumes in control group (*red*) and RT group (*black*) in response to RT (p<0.01; SD shown). (**c**) Fold change in the abundancy of the subpopulations ***b*** and ***f*** as compared to untreated tumors. A significant expansion due to RT in subpopulation ***b*** harboring Her2^+^/EGFR^+^ and subpopulation ***f*** harboring cMet^+^/MUC1^+^ is detected. Subpopulation size did not change significantly after tumor regrowth (day 17). SD is shown. **(d)** Mice were subcutaneously injected with 4T1 cells and treated with RT. Trastuzumab (**T**), 5 mg/kg, and Crizotinib (**C**), 25 mg/kg, were administrated IP 2d/week and by gavage 5d/week respectively from d0 (3 days prior RT) until the end of the experiment (d17). Std errors and p values are shown. **(e)** In-vivo fold changes in subpopulation ***b*** and ***f*** abundance showed optimal reduction when **T** and **C** were used in combination with RT. These results were consistent 6 days and 12 days after RT.

CSSS-based analysis of the tumors 6 days post-RT, when an initial shrinkage of tumors was observed (**Fig. 6b**), revealed an expansion of subpopulations **b** and **f** (**Fig. 6c**). Moreover, 12 days post-RT, when the tumors started growing again (**Fig. 6b**) we could still detect these expanded subpopulations (**Fig. 6c**).

Inhibition of both Her2 and cMet proteins significantly sensitized the tumors to RT (**Fig. 6d**). The combined treatment brought about shrinkage of the tumors and prevented development of resistance to RT (**Fig. 6d**, see the green arrow). The effect of RT plus the combined targeted therapy was highly synergistic in contrast to the effect of the two targeted drugs without RT, or RT treatment alone. Furthermore, the addition of the targeted drug combination (T+C) prior to RT brought about significant reduction in the size of subpopulations **b** and **f** (**Fig. 6e)**. No other subpopulation expanded following treatment.

### Targeting Her2 and cMet to sensitize human cell lines and patient-derived TNBC to RT

To validate that the phenomenon of the expansion of Her2+ and cMet+ cellular subpopulations is not limited to TNBC mouse models, we utilized TNBC MDA-MB-231 and MDA-MB-468 human-derived cell lines, and TNBC patient-derived cells (BR45).

Inhibition of cell growth, observed in all cell types 6 days post-RT, was followed by significant regrowth of the cells 12 days post-RT (**Fig. 7a**). Subpopulations **b** and **f**, which expanded 6 days post-RT in all cell types, maintained their size or expanded following cellular regrowth, 12 days post-RT (**Fig. 7b**).

**Figure 7.**
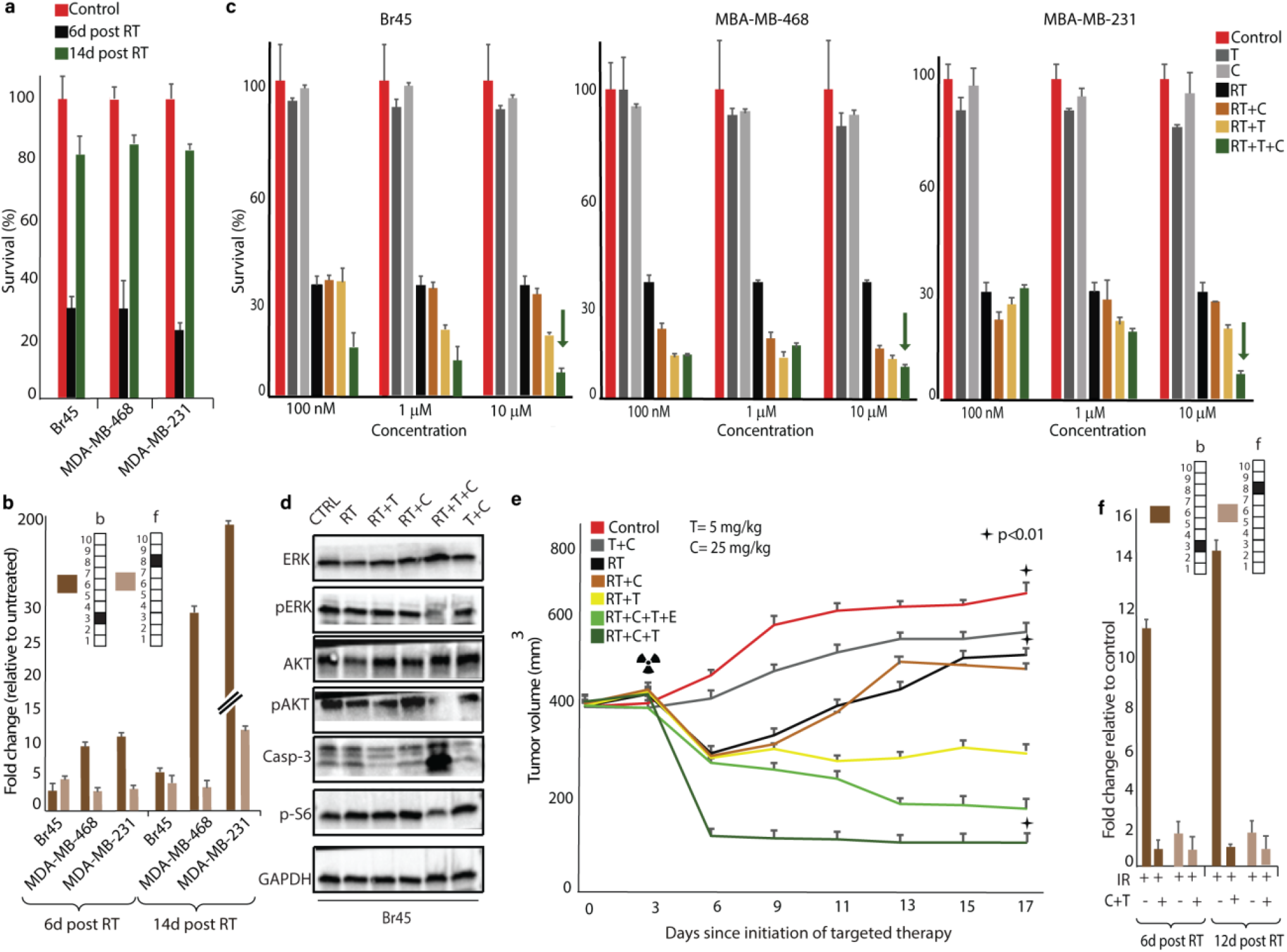
Inhibition of expanded subpopulations sensitizes human TNBC and BR45 PDX to RT. **(a)** Survival assays show a ~ 30% cell survival rate 6 days post RT, with TNBC regrowth to ~80-90% confluency 14 days post RT. **(b)** Fold changes in the abundance of subpopulations ***b*** and ***f*** compared to untreated cells. These subpopulations either remained unchanged or expanded following tumor regrowth. **(c)** Survival rates of Br45, MD-468 and MD-231 cells in response to Trastuzumab (T)+Crizotinib (C), Trastuzumab (T)+Erlotinib (E), RT, RT+T, RT+C, RT+T+E and RT+T+C 6 days post RT. Cellular drug treatment began 2 days prior to RT and was continued until the end of the experiment. **(d)** Downstream Her2 and cMet signaling protein alterations following different treatments. **C**+**T** combined with radiation induced higher levels of cleaved caspase-3, compared to irradiation alone and irradiation with either **C** or **T** alone or C+T combined. **C**+**T** administration prior to RT induced the downregulation of pAKT, pERK and p-S6 levels (**Fig. 3g**). **(e) C**+**T** sensitized TNBC response to RT in BR45 PDX in-vivo. BR45 tissues were transplanted orthotopically into 60 NSG mice treated with brachytherapy on days 3 and 5 with 12 Gy and 10 Gy respectively. Drugs were administrated from d0 (3 days prior to RT) until the end of the experiment (d17). Std errors are shown. **(f) In-vivo** fold changes in the abundance of subpopulations ***b*** and ***f*** in response to **T** and **C** use. For (**a**), (**b**) and (**c**) SD is shown.

Combined anti-Her2 and anti-cMet pretreatment sensitized all 3 types of human TNBC cells to RT (**Fig. 7c**). Each drug alone had a significantly smaller effect on cellular survival when compard to the combination of both drugs together with RT (**Fig. 7c**, see green arrow). Moreover, depletion of the downstream signaling pathways to Her2 and cMet as well as induction of cleaved caspase 3 were observed when the cells were pretreated with anti-Her2 and anti-cMET inhibitors 1 day prior to RT (**Fig. 7d**).

Using patient-derived TNBC BR45 cells grown in PDX models, we demonstrated that irradiated BR45 TNBC developed resistance to RT in a short period of time (regrowth of the tumors was detected 6 days post-RT; **Fig. 7e**, see black curve). Pretreatment of the mice with each drug alone prior RT demonstrated a small inhibitory effect on tumor growth (**Fig. 7e**). Pretreatment of the mice with the combination of both drugs however, showed significant synergistic effects with RT, bringing about significant shrinkage of the tumor and preventing the development of resistance (**Fig. 7e**, dark green curve).

Adding erlotinib (an EGFR inhibitor), which according to our algorithms was not expected to significantly influence tumor growth, did not improve the results of the Trastuzumab + Crizotinib + RT treatment combination (**Fig. 7e**).

Subpopulations **b** and **f** were reduced when the targeted drug combination (T+C) was applied prior to RT (**Fig. 7f, Fig. S5**). These results suggest that CSSS-based single cell resolution of the plasticity of TNBC in response to RT provides guidance on how effective targeted drug combinations should be designed in order to overcome RT resistance.

## Conclusions

Integrating radiobiological and biological knowledge into more efficient treatment strategies has been a major goal in the past decade. Cancer researchers and radio-oncologists are searching for new potential protein biomarkers to develop a strategy to predict and enhance tumor response to RT^40^. Although a plasticity of TNBC phenotypes in response to RT, such as a switch from a HER2- to HER2+ cellular population, has been previously detected ^13^, a strategy to exploit this plasticity and provide successful treatment is still lacking.

In this study we provided a novel framework for the resolution of intra-tumor cellular heterogeneity of aggressive TNBC. High-throughput single cell protein data was analyzed using information-theoretic surprisal analysis^25^. The analysis resolved unbalanced protein subnetworks in the tumor mass^25^, which were further assigned to single cells. Each cell was assigned a cell-specific signaling signature (CSSS), composed of a set of altered subnetworks. Cells sharing the same CSSS, were considered a subpopulation.

The demonstrated strategy not only resolved the overexpressed biomarkers or altered protein-protein correlation networks in the tumor in response to RT treatment, but also mapped single cell signaling signatures within the tumor mass. This information enabled resolution of the distinct cellular subpopulations, information that is critical for accurate treatment design.

Our analysis requires only one tissue/sample to elucidate the perturbed networks operating in each tumor. The large number of single cells analyzed (>50,000/sample) is what provides the high resolution of tumor heterogeneity. This is in contrast to bulk analysis that requires large datasets comparing multiple tissues in order to reveal the altered networks with high resolution in each patient^17,24^. Furthermore, our analysis efficiently identifies small cellular subpopulations, which are likely to be missed in bulk analyses.

Using the CSSS strategy we demonstrated that two distinct cellular subpopulations, harboring altered subnetworks with induced Her2 and cMet proteins, respectively, expanded in tumors in response to RT. Using in-vitro and in-vivo murine models, human cell lines and patient-derived TNBC, we showed that efficient sensitization of aggressive TNBC to RT could be achieved only when Her2 and cMet proteins were inhibited simultaneously.

Despite the fact that the in-vivo follow-up was only up to three weeks, the results demonstrated a significant synergistic effect in tumor response to RT and combined targeted therapy, compared to RT alone. While RT-treated tumors developed resistance, the tumors pre-treated with Her2 and cMet inhibitors demonstrated durable remission. In an extended follow-up period there is a chance that other minor sub-populations may arise, which were not seen during the three weeks. In a clinical setting a longer patient-specific follow-up might provide additional data for a more accurate treatment plan.

In summary, we demonstrate a novel approach to resolve in depth intra-tumor heterogeneity at the single cell level. This strategy provides an essential step towards the accurate design of targeted drug combinations for tumors changing phenotypes in response to RT. Elucidation and detailed analysis of TNBC plasticity allows for the sensitization of tumors to RT. Importantly, this approach allows for the mapping of distinct cellular subpopulations in a single tumor, without the need to be compared to and analyzed relative to other tumors, such as in the case of bulk analyses. The value of the proposed strategy will increase alongside the continued development of single-cell and mass cytometry techniques, which will allow for the simultaneous detecting of dozens of parameters^41^ in statistically significant numbers (>50000-1,000,000) of single cells obtained from a single tumor.

## Materials and Methods

### Cell lines and culture

Murine 4T1 cells, mimicking stage IV of TNBC model were a kind gift from Dr. Zvi Granot (Faculty of Medicine, Hebrew University of Jerusalem). Human TNBC cells MDA-MB-468 and MDA-MB-231 were acquired from ATCC and authenticated by the Genomic Center of the Technion Institute (Haifa). PDX human derived xenograft BR45 were obtained from the Oncology Department at Hadassah –Jerusalem Medical Center with prior written informed consent.

4T1 cells were routinely maintained in Dulbecco’s modified Eagle’s medium (DMEM) MDA-MB-231 and MDA-MB-468 were maintained in RPMI-1640 medium; supplemented with 10% FBS, 4 mM L-glutamine, 100 U/mL Penicillin and 100 μg/mL Streptomycin. All media and supplements were from Biological Industries, Israel. All cell lines were maintained at 37 °C in 5% CO2. Cells were checked on a routine basis for the absence of mycoplasma contamination. Irradiation of parental cells: Cells were treated by single-dose radiation with 5, 10, and 15 Gy doses of γ-rays of ^**60**^Co by a radiotherapy unit (gamma cell 220) at a dose rate of 1.5 Gy/min. See Supplementary Methods for more details.

### Murine models

#### Syngeneic models

2.0×10^**5**^ 4T1 cells were inoculated subcutaneously on 6-7 week old female Balb/c mice.

#### Alogeneic model

BR45 tumors were induced in NSG mice either by injecting 4.0×10^**6**^ cells orthopedically or by subcutaneously transplanting xenografts.

After reaching the desired tumor volume (80-100mm^3^), mice were randomly grouped to approximately 8-10 animals per cage and treatment was initialized. Tumor sizes were routinely measured with an electronic calliper every two days and their volumes were obtained using the formula V = (W (2) × L)/2. Mice were kept under conventional pathogen-free conditions. All in-vivo experiments were performed with the approval of the Hebrew University of Jerusalem IACUC. See Supplementary Methods for more details.

### In-vivo treatments

#### High dose rate (HDR) brachytherapy

Tumors were irradiated by applying a brachytherapy afterloader (GammaMed™ HDR, Iridium 192). 12 Gy was applied on two alternative days. The treatment field was designed using MRI imaging to deliver optimal radiation doses to the targeted tumors and limit exposure to surround organs at risk.

#### Targeted inhibitors

Trastuzumab (trastuzumab; Her2 inhibitor) was purchased fromTeva Pharmaceutical Industries Ltd. Crizotinib (cMet inhibitor, #12087-50) and Erlotinib (#10483-1) were purchased from Cayman Chemical. (See Suppl. Methods for doses and regimens). The treatment was based on the prediction by surprisal analysis and in-vitro validation.

### Flow Cytometry

#### Antibodies

The following fluorescently tagged antibodies, were obtained from BioLegend, Inc.: EpCAM (9C4/G8.8), CD45 (2D1/104), CD31(WM95/390), CD140a (16A1/APA5), CD44 (IM7), E-Cadherin (DECMA-1), EGFR (AY13), CD24 (M1/69), CD24 (ML5), KIT (ACK2/104D2), CD133 (315-2C11/clone7), PD-L1 (10F.9G2/29E.2A3). ERBB2 / Her2 (5J297) was obtained from LifeSpan BioScience. Anti-MUC1 Polyclonal Antibody and Anti-Met Polyclonal Antibody were both obtained from Bioss Antibodies Inc. (See Table S5).

#### Preparation of single cell suspensions and flow cytometry analysis

Following mouse euthanization, tumors were resected and mechanically disrupted to generate a single-cell suspension. Red blood cells were lysed (15mM NH4Cl + 10mM KHCO3 for 5 min at R.T.) and CD16/32 antibody was used to block the endogenous FC (#101301, Biolegend). Cells were analysed using a BD FACS LSR Fortessa. Compensation control was done using UltraComp eBeads (#01-2222-41, ThermoFisher). 50,000 cells were profiled for each sample.

Preliminary data analysis was done using FlowJo VX software. The output data were extracted into an excel file in which each row represented a single cell and each column showed the intensity of each assayed protein (FCS Extract 1.02 software). For more detailes see Supplementary Methods.

### Western blot analysis

Cell pellets were lysed with a 20% SDS buffer. The protein content of each lysate was determined with a Pierce BCA Protein Assay Kit (#23225, ThermoFisher). Equal protein aliquots were subjected to SDS-PAGE (Criterion Stain Free,4-15 % acrylamide, BIO-RAD) under reducing conditions and proteins were transferred to a nitrocellulose membrane. (Millipore). Membranes were blocked with 5% non-fat milk for 1 hour at R.T. and probed with the appropriate antibody (Supplementary Methods), followed by horseradish peroxidase-conjugated secondary antibody (#123449, Jackson ImmunoResearch) and a chemiluminescent substrate (ECL #170-5061, Bio-Rad).

### Survival assay

Cells were seeded at 70% confluency and treated as required for different time points. Cells were washed with PBS and fixed with 4% PFA for 10 min. at R.T. The fixed cells were stained with Methylene Blue (MB) for 1 hour at R.T., washed and air dried overnight. The dye was extracted with 0.1M HCl for 1 hour at R.T. Absorbance was read at 630 nm.

### MTT assay

Cells were seeded and treated as indicated in a 96 well plate for 6days. Cell viability was checked using MTT assay kit (#ab211091, Abcam). Equal volumes of MTT solution and culture media were added to each well and incubated for 3 hours at 37 *C. MTT solvent was added to each well, and then the plate was covered with aluminum foil and put on the orbital shaker for 15 minutes. Absorbance was read at 590nm following 1 hour.

### Immunofluorescence

Cells were grown on coverslips in six-well plates to reach 70% confluency by the next day, then fixed and permeabilized with cold absolute methanol. Afterwards, they were blocked with CAS blocker (cat. no. ZY-008120) and washed 3 times for 5 minutes with PBS, then stained with primary antibodies as follows: Purified anti-mouse/human Ki-67 (BLG-151202), Rabbit Anti-Met (c Met) Polyclonal Antibody (BS-0668R), Neu (F-11) SC-7301. After washing 3 times with PBS for 5 minutes, cells were stained with secondary antibodies for 1 hr at room temperature in the dark to visualize the aforementioned primary antibodies. Secondary antibodies conjugated to fluorophores were as follows: Goat anti-rat IgG H&L conjugated with Alexa Fluor 647 (1:400) (cat. no. 712605153), Affini-pure Goat anti-mouse IgG (H+L) conjugated with Alexa Fluor 488 (1:150) (cat. no. 115545003), and Affini-pure Goat anti-Rabbit IgG (H+L) conjugated with Alexa Fluor 488 (1:150) (cat. no. 111545003). All secondary antibodies were purchased from Jackson ImmunoResearch. After washing 3 times with PBS, cell slides were mounted using fluorogel III mixed with DAPI (Bar Naor, cat. no. 17985-01) to stain the nuclei. A Spinning Disk Confocal microscope was used to visualize the expression of biomarkers of interest. The analysis was done with NIS elements software.

### Single cell data analysis

The analysis is composed of two steps: First - single cell surprisal analysis (SA) is utilized to identify unbalanced processes in the cellular population as previously described^25^. Briefly, the data matrix obtained from the flow cytometry analysis, in which columns are expression levels of the tested proteins and rows are single cells, is used as an input for surprisal analysis (calculations are performed in MATLAB). The analysis is based on the premise that all biological systems reach a state of minimal free energy under standard temperature and pressure given the existing environmental and genomic constraints. The analysis identifies protein expression levels at the reference state, and the deviations in those levels due to the existing constraints. RT treatment imposes a constraint but more than one constraint may be identified in the system. Each constraint is associated with an altered protein subnetwork (= unbalanced biological process) that deviates the system from the reference state and causes the coordinated deviations of a subset of proteins from their steady state expression level.

In heterogeneous tissues many processes occur through the actions of individual cells. Thus the analysis was implemented independently for each measured cell. The levels of different proteins for each cell at each time point t are represented as Equation 1:

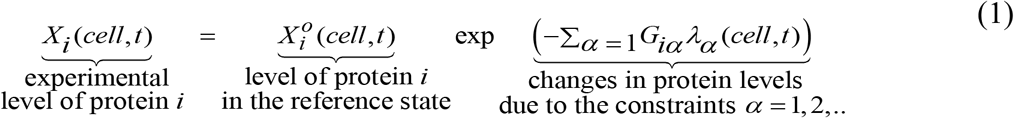

Here,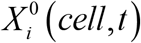 (*cell,t*) is the expected expression level of protein *i* at the reference state in a measured cell at the time point *t*. The exponential term in Equation 1 represents the deviation from the reference value due to the constraints, including those imposed by Irradiation. *G*_*iα*_ are weights of protein *i* in the unbalanced processes *α* = 1, 2.. Proteins deviating in a similar manner from the steady state are grouped into unbalanced processes (Figure 3). *λ* _*a*_ (*cell,t*) is the amplitude of an unbalanced processes *α* = 1, 2.. in a cell *i* at time point *t*. (Example for *G*_*iα*_ values, as calculated for 4T1 models is presented in Table S4. Several unbalanced processes can be found in the system, however not all processes are active in all cells.

Second step: To further map distinct subpopulations within the entire cellular population, where all the cells sharing the same set of unbalanced processes, or CSSS, are grouped into subpopulations (Figure 3). Each CSSS is transformed into a barcode for the simplicity of calculations and representation.

#### Barcode calculations

The output lambda file from the surprisal analysis is then used as an input file for the Python script in order to obtain a specific barcode for each single cell in which a certain unbalanced process is active/inactive. Briefly, *λ*_*a*_(*cell,t*) values are sorted and plotted as sigmoid plots in each process. Only *λ*_*a*_(*cell,t*) values located on the tails of the sorted distributions are considered and used further for the barcode calculations (Figure S4). Several processes may be active in each cell, as amplitudes, *λ*_*a*_(*cell,t*), of several processes may be significant in each cell. See Supplementary Methods section for more details.

### Bulk analysis of gene expression data

The expression level of each transcript is decomposed by surprisal analysis due to environmental or genomic constraints present in the system ^14,24^. Any genetic defect or epigenetic perturbation can impose a constraint that alters a part of the gene expression network structure in the system, which in turn causes specific group of transcripts (=subnetwork) to undergo coordinated changes in their expression levels. This group of co-varying transcripts is defined as an unbalanced process. Each of the altered transcripts can be involved in several unbalanced processes due to the non-linearity of biological networks. To deconstruct gene expression levels into the levels at the steady state and deviation thereof, the following equation is utilized: In 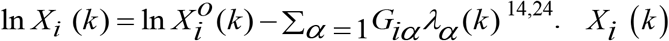 is the actual, experimentally measured expression level of the transcript i in a cancer sample k. 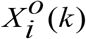 are the expression levels at the steady state. In cases where 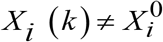, we assume that the expression level of transcript I was altered due to constraints that operate inn the system. The analysis uncovers the complete set of constraints and the assosiated unbalanced processes.

The unbalanced processes are indexed by α = 1,2,3. Several unbalanced processes may operate in each tumor. The term ∑_*α* = 1_*G*_*iα*_*λ*_*α*_(*k*) represents the sum of deviations in expression level of protein i due to the various constraints, or unbalanced processes that exist in the sample. The *λ*_*α*_(*k*) values, denote the amplitude of each unbalanced process, in every sample *k*, i.e. the extent of the participation of each unbalanced process α, in every sample/tumor *k*. The amplitude, *λ* _*α*_ (*k*) (**Table S2**), determines whether process α is active in the sample k, and to what extent. A detailed description of the surprisal analysis process can be found in ^14,17,24^. (2) The *G*_*iα*_ values (**Table S1**), denoting the extent of participation of each individual protein *i* in the specific unbalanced process, *α*.

#### Determination of the number of significant unbalanced processes

As described previously ^24,37^.

#### Meaning of the negative and positive signs in the analysis

The term *G*_*iα*_ denotes the degree of participation of the transcript *i* in the unbalanced process α, and its sign indicates the correlation or anti-correlation between transcripts in the same process. For example, in a certain process α, transcripts can be assigned the values: G_transcript1,α_ = −0.013, G_transcript2,α_ = 0.02, and G_transcript3,α_ = 0.0003, indicating that this process altered transcripts 1 and 2 in opposite directions (i.e. transcript 1 is upregulated and transcript 2 is downregulated, or vice versa due to the process α), while not affecting transcript 3.

Importantly, not all processes are active in all samples. The term *λ*_*α*_*(k)* represents the importance of the unbalanced process *α* in the tumor *k*. Its sign indicates the correlation or anti-correlation between the same processes in different tumors. For example, if process α is assigned the values: *λ*_*α*_*(1)* = 3.1, *λ*_*α*_*(2)* = 0.02, and *λ*_*α*_*(5)* = 2.5, it means that this process influences the tumors of the patients indexed 1 and 2 in the same direction, while it is inactive in patient 5.

To calculate an induction or reduction due to process *a*, a product *G*_*iα*_*λ*_*α*_*(k)* is computed.

To provide a biological interpretation of each unbalanced process only those transcripts that were located on the tails (Table S1) were included in the analysis. The classification of the transripts into biological categories was performed using David database and presented in Table S1.

## Acknowledgements

The project was funded by the Israel Science Foundation (ISF).

We acknowledge Science Training Encouraging Peace (STEP) for partial funding of this research, and Efrat Flashner-Abramson (H.A’s STEP partner) for helpful input.

## Author contributions

H.A., A.M.R., A.M. and N.K.-B. designed research; H.A., A.M.R., A.M., S.V., S.S., K.S., A.M. and N.K.-B. performed research; S.V., E.F.-A., S.S.,T.P.Y., A.G.,Z.G., I.B.P amd A.M. contributed experimental/analytical tools. H.A., A.M.R., S.V. and N.K.-B. analyzed data; H.A., E.F.A., A.M., K.S. and N.K.-B wrote the paper with contributions from all authors.

## The authors declare no conflict of interest

### Data availability

The data supporting the findings of this study are available in this paper or the <u>Supplementary Information.</u>Any other raw data that support this study are available from the corresponding author upon request.

