## Supplementary Information for "Mapping cellular subpopulations within triple negative breast cancer tumors provides a tool for cancer sensitization to radiotherapy"

### Supplementary Methods

#### Single Cell Surprisal analysis.

Surprisal analysis is a thermodynamic-based information-theoretical approach [1–3]. The analysis is based on the premise that biological systems reach a balanced state when the system is free of constraints [4–6]. However, under the influence of environmental and genomic constraints, the system is prevented from reaching the state of minimal free energy, and instead reaches a state which is higher in free energy (in biological systems, which are normally under constant temperature and constant pressure, minimal free energy equals maximal entropy).

For example, if the system under study is a living cell, an environmental constraint can be exposure to a drug, which inflicts a change in protein concentrations and activities in the cell. The system can be influenced by genomic constraints as well, such as genomic mutations that in turn affect protein function, often eliciting alteration of specific signaling pathways to oppose the functions of the damaged protein.

Surprisal analysis can take as input the expression levels of various macromolecules, e.g. genes, transcripts, or proteins. However, be it environmental or genomic alterations, it is the proteins that execute the main functions of a cell, and therefore we base our analysis on proteomic data. The varying forces, or constraints, that act upon living cells ultimately manifest as alterations in the cellular protein network. Each constraint induces a change in a specific part of the protein network in the cells. The subnetwork that is altered due to the specific constraint is termed an *unbalanced process*. System can be influenced by several constraints thus leading to the emergence of several unbalanced processes. When tumor cells are characterized, the specific set of unbalanced processes can be active in a cell. This is what constitutes the cell-specific signaling signature.

In heterogeneous tissues many processes can occur through the actions of individual cells. Thus the analysis was implemented independently for each measured cell. The levels of different proteins for each cell at each time point *t* are represented as Equation 1:

$$\underbrace{X_i(cell,t)}_{\substack{\text{experimental}\\ \text{level of protein } i}} = \underbrace{X_i^o(cell,t)}_{\substack{\text{level of protein } i\\ \text{in the reference state}}} \exp \underbrace{\left(-\Sigma_{\alpha=1} G_{i\alpha}\lambda_\alpha(cell,t)\right)}_{\substack{\text{changes in protein levels}\\ \text{due to the constraints } \alpha=1,2,..}} \qquad (1)$$

For every protein, *i*, surprisal analysis calculates the distribution of the expression levels at the reference state: $X_i^0$. This term was shown to be constant, i.e. is independent of time and of the actual state of the system [7–10]. In terms of information theory, $X_i^0$ represents the state of maximal entropy, or minimal information.

Here, $X_i^0(cell,t)$ the expected expression level of a protein *i* at the reference state in a measured cell at the time point *t*. The exponential term in Equation 1 represents the deviation from the reference value due to the constraints, including those imposed by Irradiation. $G_{i\alpha}$ are weights (the degree of participation) of a protein *i* in the unbalanced processes $\alpha = 1, 2..$ Proteins deviating in a similar manner from the steady state are grouped into unbalanced processes (Figure 2 and 3). $\lambda_a(cell,t)$ is an amplitude of an unbalanced processes $\alpha = 1, 2..$ in a cell *i* at time point *t*. (Example for $G_{i\alpha}$ values , as calculated for 4T1 models is presented in Table S4. Figure S3 represents

$\lambda_3(cell,t)$ values for process 3. Several unbalanced processes can be found in the system, however not all processes are active in all cells.

$G_{i\alpha}$ sign indicates the correlation or anti-correlation between proteins in the same process. For example, in a certain process $\alpha$, proteins can be assigned the values: $G_{\text{protein 1},\alpha} = -0.50$, $G_{\text{protein 2},\alpha} = 0.44$, and $G_{\text{protein 3},\alpha} = 0.00$, indicating that this process altered expression levels of the proteins 1 and 2 in opposite directions while not affecting protein 3. Each protein can take part in a number of unbalanced processes at once.

Note that in order to define upregulation or downregulation in protein expression levels due to a specific process $\alpha$ the product $G_{i\alpha}\lambda_{\alpha}(cell,t)$ is calculated.

Importantly, not all processes are active in all cells. The term $\lambda_{\alpha}(cell,t)$ represents the importance of the unbalanced process $\alpha$ in *cell*. Its sign indicates the correlation or anti-correlation between the same processes in different cells. For example, if the process $\alpha$ is assigned the values: $\lambda_{\alpha}(1) = 3.1$, $\lambda_{\alpha}(2) = 0.0$, and $\lambda_{\alpha}(3) = 2.5$, it means that this process influences the cells indexed 1 and 3 in the same direction, while it is inactive in cell 2.

**Generation of functional networks.**

Figure S2 complements Fig 4f in the main text and represents other functional networks active in the system. The goal was to generate unbalanced processes composed of proteins with significant $G_{i\alpha}$ values. Functional connections between the proteins in each unbalanced process are based on STRING database.

**Calculation of barcodes.**

The barcodes presented in Figure 4g were generated using a python script (written with the assistance of Mr. Jonathan Abramson). In this script, for each process α, $\lambda_\alpha(cell)$ $\alpha = 1, 2, 3, \ldots, 10$) values were normalized as follows: If, e.g.; $\lambda_\alpha(cell) > 0.5$ (and is therefore significant according to calculation of threshold (limit) values) then it was normalized to 1; if $\lambda_\alpha(cell) < -0.5$ (significant according to threshold values as well) then it was normalized to -1; and if $-0.5 < \lambda_\alpha(cell) < 0.5$ then it was normalized to 0. These thresholds (limits) were obtained after $\lambda_a(cell)$ values were sorted according to their values, and only cells with significant $\lambda_a(cell)$ values were considered to possess an unbalanced process α. Only $\lambda_a(cell)$ values located on the tails of the sorted distributions are considered significant. For more details see Figure S4.

**Irradiation of parental cells.**

4T1, MDA-MB-231, MDA-MB-468 and Br45 cells were trypsinized and plated to reach optimal confluences next day by (70-80) % before irradiation treatment. The next day, 4T1 cells were irradiated using (5 and 15)Gy of γ-rays. Radiation doses were selected based on calibration experiments in which the survival rates after irradiation ranged from (40-50)% of the cells. MDA-MB-231 and BR45 cells were treated with 10 Gy with the exception of MDA-MB-468 cells which were treated with 5 Gy. Afterwards, cells were grown under normal conditions for 24h, 48h and 6 days. At each indicated time point, cells were detached from the flask using *Acutase* and fixed using 2% paraformaldehyde for 30 min in ice. Labelling procedure of each condition performed on the day of the flow cytometry analysis as mentioned below.

**Mouse models and tumor inoculation.**

4T1 mouse breast carcinoma mimics stage IV triple negative breast cancer in human. It is highly metastatic to lungs, lymph nodes, liver and bones after implantation in the mammary fat pad of immune-competent Balb/c mice. Primary tumors were harvested after euthanizing female mice using carbon dioxide ($CO_2$) inhalation. Time elapsed from tumor inoculation varied from 2 weeks to 1 month. Human Br45 was used as PDX model, tumors were harvested 2 months after surgically transplanting xenografts on NSG-NOD mouse model or after 4 months from injecting Br45 cells orthopedically.

**In-vivo treatments.**

Targeted inhibitors:

Drugs were given usually by IP or gavage depending on the drug. Herceptin was given IP twice a week with a concentration of 5mg/kg, the vehicle was 200ul sterile saline. Crizotinib and erlotinib were given by gavage with a concentration of 25mg/kg and 12.5mg/kg respectively, five days a week. The vehicle used was hydroxypropyl methylcellulose with 0.2% tween. Mice were treated for 3 weeks. During this period the tumor volumes were measured regularly to observe the action of the drug.

**Flow Cytometry method.**

**Antibodies:** Detailed information about all the FACS antibodies are found in Table S5.

**Preparation of single mammary tumor cell suspensions:**

Dissecting of mammary gland tumor from mice: Small incision is made on the level of the skin and the tumor mass is gently separate from the conjunctive tissue using a sharp blade.

Preparing of single cell suspensions using mechanical method: The harvested tumors were washed twice with PBS at RT and minced thoroughly. Then the masses are smashed gently with the back of 10 ml plastic syringe to mechanically digest them. After being placed in the stir apparatus for 15-20 min, the tumor/PBS buffer mixture is to be strained through a 70µm cell strainer and then centrifuged for 5 min at 3000 rcf.

**Red blood cells lysis:** To lyse RBCs from the freshly harvested tumors, cells were re-suspended in 10 ml RBC lysis buffer (0.8 g $NH_4CL$ + 0.1 g $KHCO_3$ in 100 ml DDW) for 5 min. at RT. To stop the reaction of the buffer, 30 ml of DMEM with 10% FBS was added and then the mixture was centrifuged for 5 min at 3000 rcf to get rid of the lysis buffer.

**Fixation of cells:** Samples were fixed and permeabilized with 2% PFA (#15710, EMS ) for 30min in ice.

**<u>Blocking of Endogenous Fc</u>**: 1.5 ml centrifuge tube, each contains $0.8\times10^6$ cells were incubated for 30 mins in ice with 50 ul of Fc blocker buffer (FACS Buffer + Fc Blocker: anti-mouse CD16/32 Antibody, Biolegend #101301, 1:50).

**<u>Labelling Procedure</u>:** Each sample was labelled with 11 fluorescently tagged Abs mixture. (Table S5). A cocktail of 3 additional Abs with the same fluorophore (PE) used as an exclusion criteria for hematopoietic (CD45), fibroblasts (CD140) and endothelial cells (CD31) to ensure that only tumor cells will be analysed later on. This criterion is not needed in case of staining parental cells which do not have tumor microenvironment. The 11 fluorescently labelled Abs are detailed in Table S5. Unstained control sample for each condition was used along with a single colour control for each Ab using UltraComp Compensation eBeads**™** according to the manufacturer's instructions for creating compensation controls. The labelling time extended to 40 min in ice in the dark. Then, samples were washed with an equal amount of flow FACS buffer, centrifuged and resuspended in 700 µl of flow FACS buffer then filtered right before reading with LSR-Fortessa Analyser into FACS tubes.

**Western blot analysis.**

<u>Antibodies</u>: Western blot antibodies were obtained from Cell Signaling Technology, Inc.: anti-phospho-Akt ( Thr308, #4056S), anti-phospho-Akt (Ser473, #9271S), anti-total-Akt (#4691S), anti-phospho-ERK1/2 Thr202/Tyr204 (#9101S), anti-total-ERK1/2 (#9102S), anti-cleaved PARP(#5625S), Cleaved Caspase-3 (Asp 175, #9661S), Phospho-S6 Ribosomal Protein (ser235/236, #2211S) (D57.2.2E) XP® Rabbit mAb. GAPDH Antibody (#32233) was obtained from Santa Cruz Biotechnology Inc.

### Supplementary Figures and legends

### Figure S1.

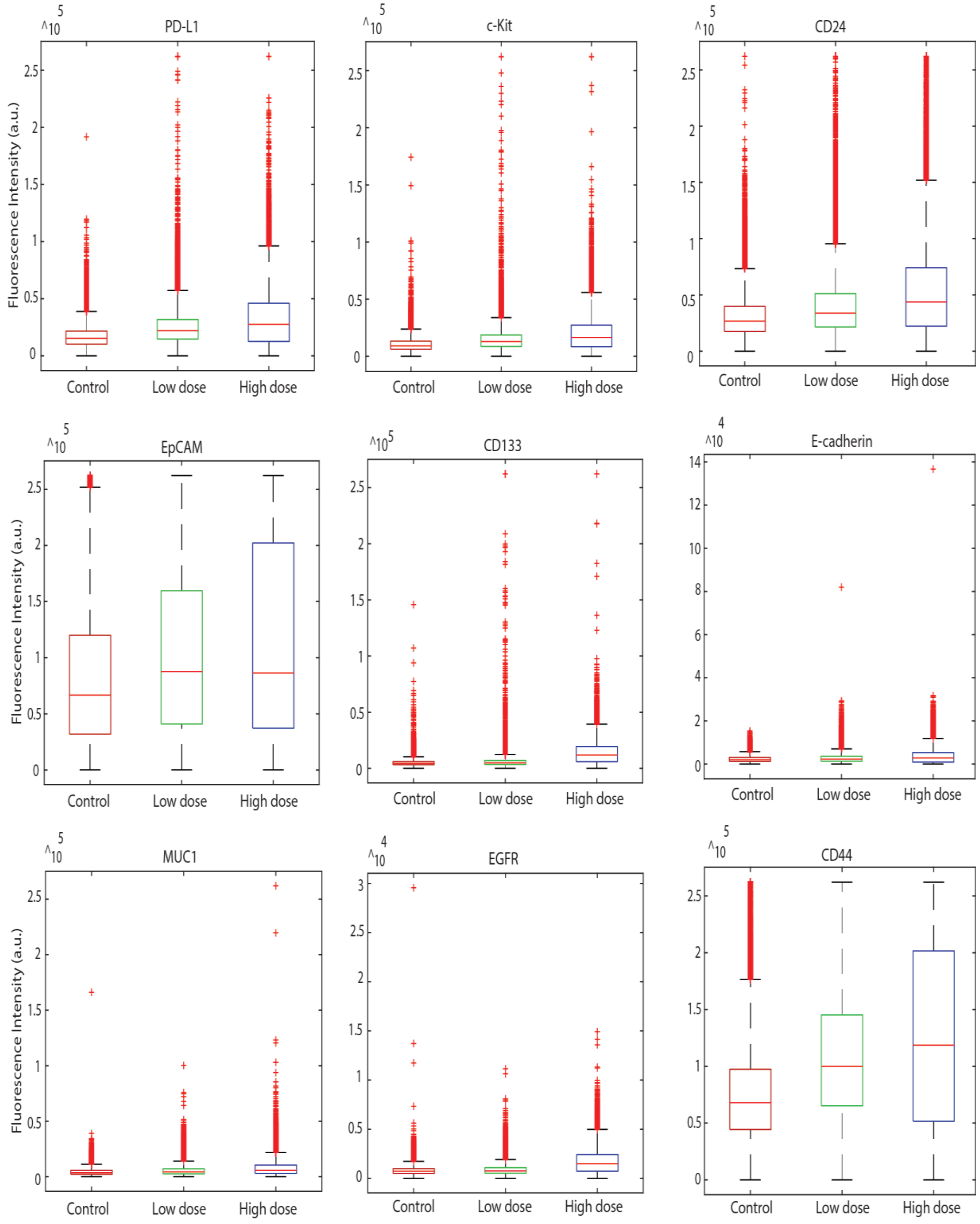

**Figure S1. Expression levels of 11 proteins, before and after irradiation in 4T1 cells.** Raw FACS data of protein expression levels in response to low dose (5Gy) or high dose (15 Gy) are shown as one-dimensional boxplots.

**Figure S2.**

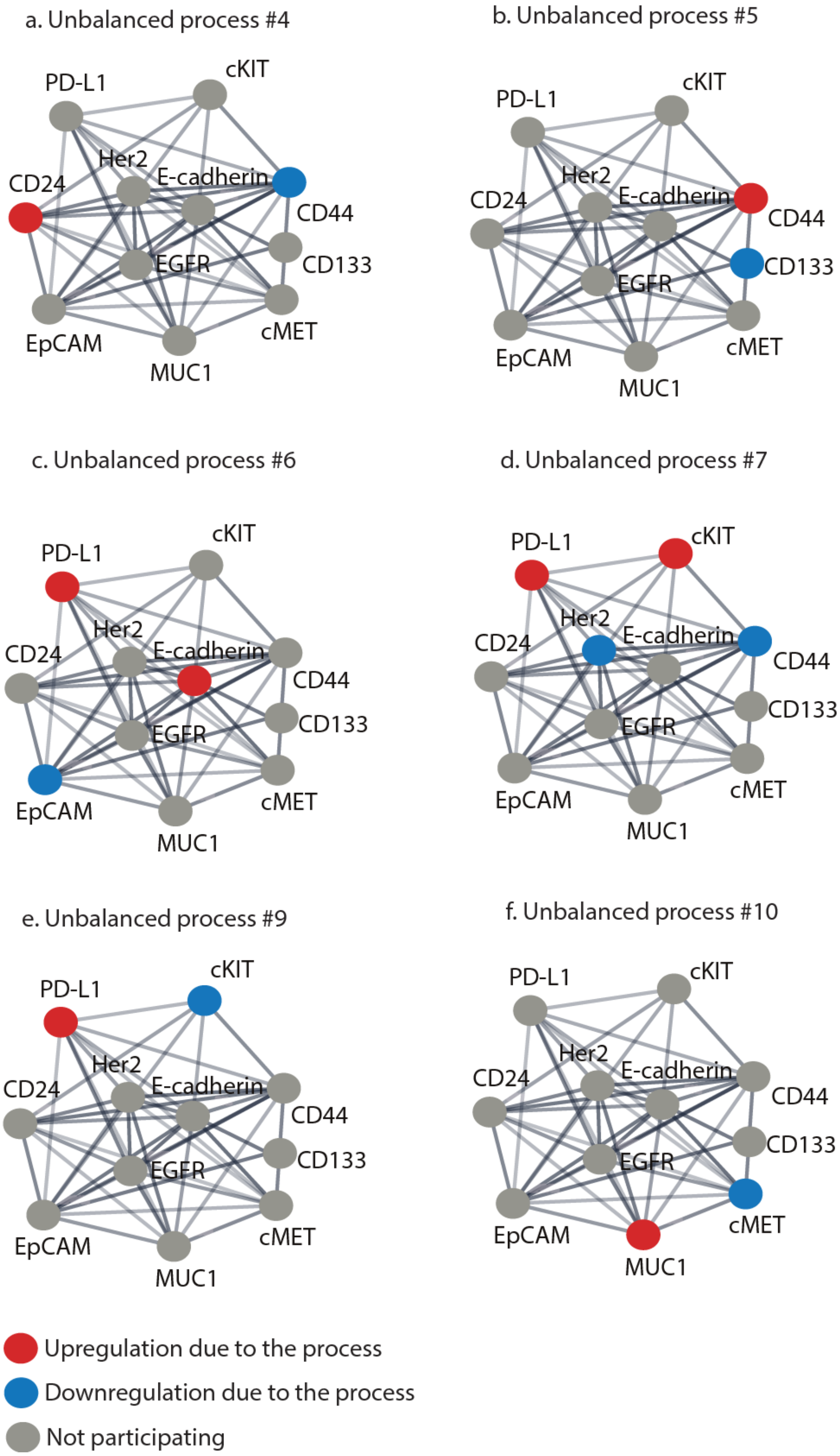
a. Unbalanced process #4
b. Unbalanced process #5
c. Unbalanced process #6
d. Unbalanced process #7
e. Unbalanced process #9
f. Unbalanced process #10
PD-L1
cKIT
Her2
CD24
E-cadherin
CD44
CD133
EGFR
EpCAM
cMET
MUC1
Upregulation due to the process
Downregulation due to the process
Not participating

**Figure S2. Unbalanced processes (=subnetworks) $\alpha = 4-10$ as identified by surprisal analysis for 4T1 irradiated cells.** (a-f) For every process $\alpha$, the proteins with significant $Gi_{\alpha}$ values were assembled into subnetworks. Upregulation and downregulation due to the process was calculated using a product $G_{i\alpha}\lambda_{\alpha}(cell,t)$ for each *cell* at a time point *t*. All proteins involved in the process were assigned functional connections using STRING database.

**Figure S3.**

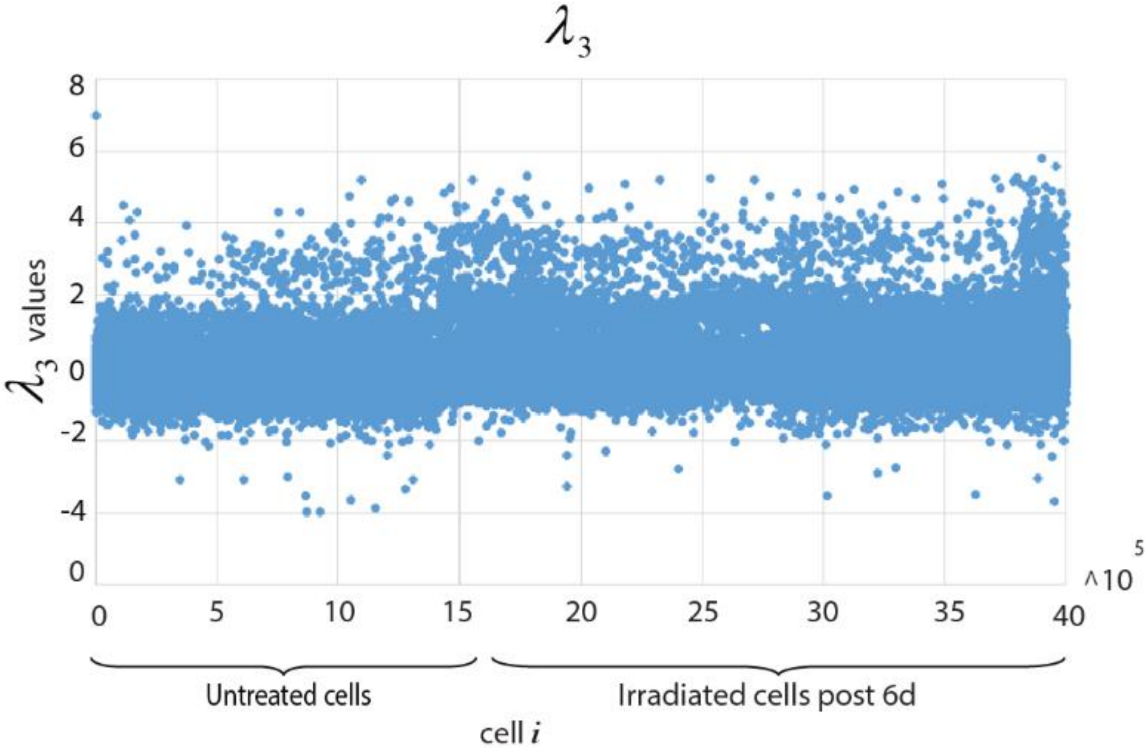

**Figure S3. Scatter plot representation of $\lambda 3(cell,t)$ values for process 3 in 4T1 irradiated cells is shown as an example.** The amplitude of unbalanced process 3 is plotted in order to follow up with the information this process is providing. The proteins that are participating in this process are Her2 and to a lesser extent EGFR. Based on the calculation of $G_{i\alpha}\lambda_{\alpha}(cell,t)$ product, the upregulated and downregulated protein expression levels are being defined. Due to this process 3, EGFR and Her2 are being altered in the same direction (EGFR is upregulated and Her2 is also upregulated or vice versa due to the process). The rest of the 11 proteins used in our panel are not affected by this process knowing that one protein can participate in a number of unbalance processes in the same time. Similar plots are built for all lambda values.

**Figure S4.**

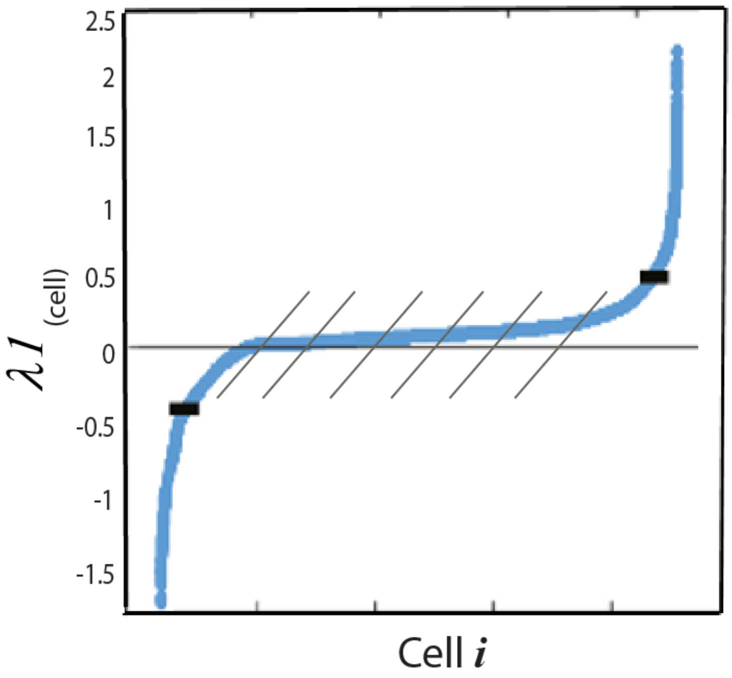

**Figure S4. Sigmoid plotting of $\lambda 1(cell)$ values to identify the thresholds (limits) of significant values posing an altered process 1.** Active processes in different cells were identified as follows: For every unbalanced process α, $\lambda_\alpha$ (cell) values were sorted according to their values, and only cells with significant $\lambda_\alpha$ (cell) values were considered to possess an unbalanced process α. This is exemplified for the process α = 1 of 4T1 cells post to RT as shown in the figure. Shown are sorted values of $\lambda_1$ (cell), which represent an amplitude of the process α = 1 in each cell. The black lines represent threshold values. In this example cells received the values $\lambda_\alpha$ (cell) > 0.5 or $\lambda_\alpha$ (cell) < -0.5 (which form the top and bottom "tails" of the distribution) for the process #1, were considered to possess the unbalanced process α = 1 and used in the python script in order to obtain the barcodes shown in Figure 4g. These values were used to calculate further the products $G_{i\alpha}\lambda_\alpha(cell,t)$ in order to build a functional subnetwork using STRING database, presented in Figure S2.

**Figure S5.**

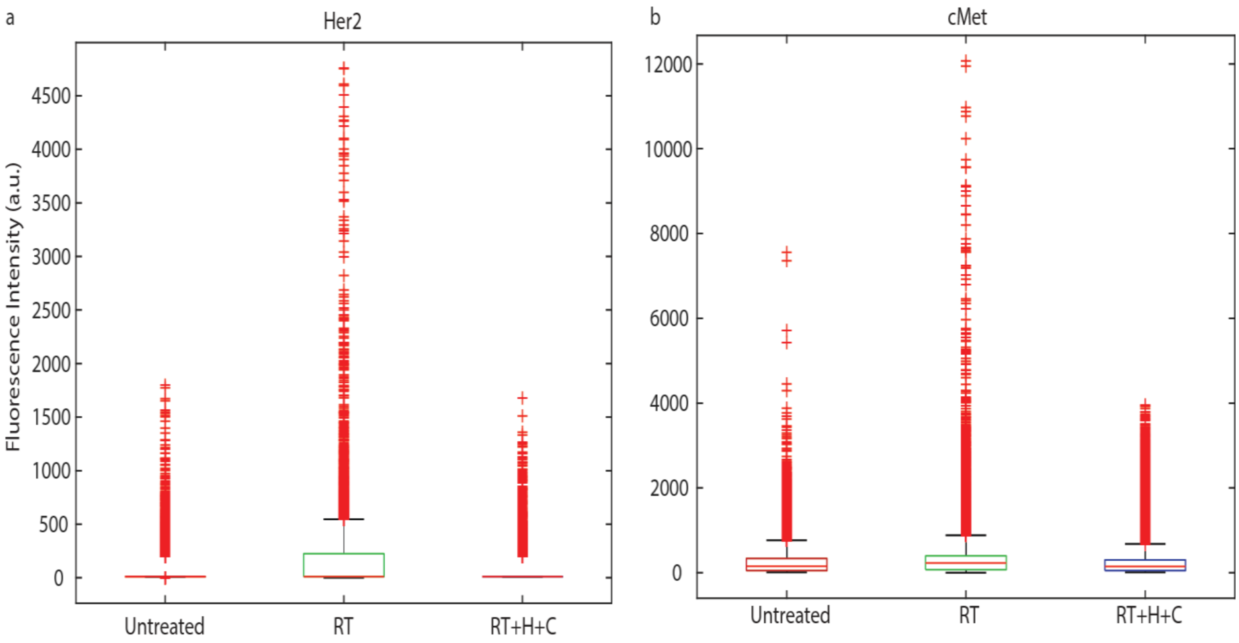

**Figure S5. In-vivo change in the expression levels of Her2 and cMet in Br45 cells.** One-dimensional boxplots show the upregulation of Her2 (a) and cMet (b) after irradiation on two alternative days with (10, 12 ) Gy respectively and the downregulation of these proteins when CSSS-predicted targeted therapy was applied.

**Supplementary Tables.**

**Table S3.**

| Process | Control | 24 hrs. post IR | 48 hrs. post IR | 6 days post IR |
|---|---|---|---|---|
| # 1 | 25% | 1.2% | 1.6% | 11.1% |
| # 2 | 18% | 14.6% | 10% | 11% |
| **# 3** | **0.3%** | **1.2%** | **2.8%** | **22%** |
| # 4 | 0.4% | 0.3% | 0.2% | 0.5% |
| # 5 | 1.5% | 0.6% | 0.5% | 1.4% |
| # 6 | 0.2% | 0.02% | 0.08% | 0.2% |
| # 7 | 0.2% | 0.6% | 0.05% | 0.3% |
| **# 8** | **0.5%** | **0.7%** | **1%** | **4%** |
| # 9 | 0.1% | 0.1% | 0.09% | 0.5% |
| # 10 | 0.1% | 0 | 0 | 0.06% |

**Table. S3 Calculating the percentages of cell subpopulation for each single unbalanced processes in 4T1 irradiated cells.** The surprisal analysis revealed 10 unbalanced processes (i.e. altered protein-protein correlation patterns resulting from 10 constraints) which occurred in the untreated/treated cells. Five of the processes, **(#1**, **#2**, **#3**, **#5** and **#8)** are appearing in at least 1% of the *treated cells*. The most abundant processes, indexed 1 and 2, appeared in 25% and 18% of the untreated cells, respectively. Processes 3 and 8, which included correlated Her2/EGFR and cMet/Muc1, correspondingly, initially demonstrated low abundancy, and appeared in 0.3% and 0.5% of the untreated cells, correspondingly. Processes 3 and 8 became more dominant 6 days post-RT.

**Table S4.**

| proteins | G1 | G2 | G3 | G4 | G5 | G6 | G7 | G8 | G9 | G10 |
|---|---|---|---|---|---|---|---|---|---|---|
| Her2 | **0.4** | -0.6 | **0.9** | 0.007 | 0.07 | -0.04 | -0.001 | -0.02 | 0.005 | -0.006 |
| EGFR | **-0.9** | 0.3 | **0.4** | 0.1 | 0.2 | -0.02 | 0.06 | -0.03 | 0.002 | -0.01 |
| EpCAM | -0.01 | **-0.4** | -0.1 | -0.2 | 0.2 | -0.4 | 0.2 | **-0.5** | -0.02 | 0.1 |
| CD24 | 0.13 | 0.04 | -0.1 | **0.8** | 0.1 | -0.1 | -0.3 | 0.02 | 0.1 | 0.003 |
| CD44 | 0.26 | **0.6** | -0.1 | **-0.4** | 0.3 | -0.1 | -0.4 | 0.04 | -0.03 | -0.01 |
| PD-L | 0.18 | 0.2 | -0.1 | 0.1 | 0.01 | **0.4** | **0.6** | -0.2 | **0.5** | 0.03 |
| cKit | 0.1 | 0.02 | -0.1 | 0.1 | 0.03 | 0.2 | **0.4** | 0.2 | **-0.8** | 0.04 |
| CD133 | -0.04 | 0.2 | 0.04 | -0.03 | **-0.8** | -0.3 | 0.02 | 0.2 | 0.02 | 0.004 |
| Ecadherin | -0.1 | -0.2 | 0.02 | -0.1 | -0.2 | **0.6** | **-0.4** | **-0.4** | -0.2 | -0.1 |
| cMet | -0.04 | -0.3 | 0.05 | -0.1 | 0.05 | -0.01 | 0.01 | **0.4** | 0.2 | **-0.7** |
| MUC1 | -0.1 | -0.3 | 0.01 | -0.1 | 0.01 | 0.1 | -0.1 | **0.5** | 0.2 | **0.6** |

**Table. S4 G values for 4T1 irradiated cells.** $G_{i\alpha}$ **are weights of a protein** $i$ **in the unbalanced processes (**$\alpha = 1, 2..$**).** Significant G values are labeled in blue and red colors. The sign of the G value indicates the correlation or anti-correlation between proteins in the same process as shown in the table. $G_{\text{Her2 }1} = 0.4$ and $G_{\text{EGFR }1} = -0.9$ indicating that in process #1 the two altered proteins are Her2 and EGFR are anti-correlated due to the process.

**Table S5. Antibodies for flow cytometry analysis.**

| Protein | Reactivity | Color | Company | Cat. # | Concentration ($\mu l/8\times10^5$ cells) |
|---|---|---|---|---|---|
| CD45 | Mouse (M) | PE | Biolegend | 103105 | 0.2 |
| CD140 | M | PE | Biolegend | 135905 | 1 |
| CD31 | M | PE | Biolegend | 102407 | 0.3 |
| CD326 – EpCAM | M | APC | Biolegend | 118213 | 1 |
| CD24 | M | BV421 | Biolegend | 101825 | 0.75 |
| CD117 – cKit | M | BV650 | Biolegend | 135125 | 1.5 |
| CD274 – PD-L1 | M | BV605 | Biolegend | 124321 | 1.5 |
| CD133 | M | PE/Dazzel594 | Biolegend | 141211 | 1.5 |
| CD45 | Human (H) | PE | Biolegend | 638512 | 0.2 |
| CD140 | H | PE | Biolegend | 323504 | 1 |
| CD31 | H | PE | Biolegend | 303405 | 0.25 |
| CD326 – EpCAM | H | APC | Biolegend | 324207 | 1 |
| CD24 | H | BV421 | Biolegend | 311121 | 0.75 |
| CD117 – cKit | H | BV650 | Biolegend | 313221 | 1.5 |
| CD274 – PD-L1 | H | BV605 | Biolegend | 329723 | 1.5 |
| CD133 | H | PE/Dazzel594 | Biolegend | 372811 | 1.5 |
| CD44 | H/M | BV510 | Biolegend | 103043 | 1.5 |
| cMet | H/M | Alexa Fluor 750 | Bioss | 0668R-A750 | 1.5 |
| MUC1 | H/M | Alexa Fluor 680 | Bioss | 1497R-A680 | 1.5 |
| Her2 | H/M | FITC | LifeSpan BioScience | C533753 | 1 |
| EGFR | H/M | Per/CP5.5 | Santa Cruz | 20 PCPC5 | 1 |
| CD324 – ECAD | H/M | PE/Cy7 | Biolegend | 147309 | 1.5 |

9. Kravchenko-Balasha N, H. Johnson, F.M. White, J.R. Heath, Levine R.D. A Thermodynamic Based Interpretation of Protein Expression Heterogeneity in Different

GBM Tumors Identifies Tumor Specific Unbalanced Processes. J Chem Phys. 2016;

10. Zadran S, Remacle F, Levine RD. miRNA and mRNA cancer signatures determined by analysis of expression levels in large cohorts of patients. Proc Natl Acad Sci U S A. 2013;110(47):19160–5.
